# Integrated omics analysis reveals reorganization of nitrogen and lipid metabolism in a toluene-degrading bacterium in the gas phase

**DOI:** 10.64898/2026.03.26.714097

**Authors:** Shori Inoue, Taisei Naobayashi, Kanako Tokiyoshi, Shogo Yoshimoto, Hiroshi Tsugawa, Katsutoshi Hori

## Abstract

Gas-phase bioprocesses, in which microbial cells are immobilized on solid carriers exposed to the gas phase, enable efficient conversion of poorly water-soluble compounds for biodegradation and bioproduction. However, operation without a surrounding bulk aqueous medium can constrain the replenishment of water and nonvolatile nutrients. The metabolic mechanisms that sustain cellular activity in these low-water environments remain largely unknown. Here, we elucidated the metabolic alterations in the highly adhesive bacterium *Acinetobacter* sp. Tol 5 degrading toluene in the gas and aqueous phases. Polyurethane carriers bearing cells from independent cultures were assigned to the gas phase or to two aqueous media, nitrogen-depleted basal salt medium and phosphate-buffered saline, so that the conditions differed only in phase and nutrient availability. In cells actively degrading toluene, integrated metabolomics, lipidomics, and transcriptomics (97 metabolites, 108 lipids, and 3,899 genes) revealed a metabolic state specific to the gas phase. Under gas-phase conditions, amino acid and nucleotide degradation was enhanced while the intracellular glutamate pool was maintained at a high level, and citrulline specifically accumulated. Membrane lipids were remodeled, and the storage lipid triacylglycerol was consumed, suggesting that it serves as an endogenous carbon and energy source and may also contribute to metabolic water production under nutrient- and water-limited gas-phase conditions. These findings highlight nitrogen metabolism and lipid mobilization as features of gas-phase adaptation, providing a rational basis for designing robust gas-phase bioprocesses and selecting suitable microbial chassis.

## 1 Introduction

Microbial bioprocesses utilizing gaseous substrates, including greenhouse gases, syngas, and volatile organic compounds (VOCs), have emerged as pivotal strategies for preventing air pollution and establishing a sustainable carbon cycle. While microbial waste gas treatment has traditionally been developed in the environmental field for pollution control (Rybarczyk et al., 2019; Sheoran et al., 2022), recent research has increasingly focused on the valorization of these gaseous substrates into fuels and value-added chemicals (Liew et al., 2022; Tan et al., 2024; Xia et al., 2024; Devi and Pakshirajan, 2025). A major bottleneck is often gas-liquid mass transfer, which is limited by the low aqueous solubility of these substrates and necessitates energy-intensive aeration and agitation (Asimakopoulos et al., 2018; Neto et al., 2024). To overcome this limitation, various reactor configurations have been proposed. For instance, hollow fiber membrane reactors and trickle bed reactors have been reported to achieve high mass transfer rates without intensive aeration or agitation (Takors et al., 2018; Neto et al., 2024). Furthermore, novel reactor designs that expose cells directly to the gas phase, such as inverse membrane bioreactors and horizontally oriented rotating packed bed reactors, have demonstrated enhanced productivity by improving the mass transfer of poorly soluble gases (Shen et al., 2017; Chen et al., 2023).

The toluene-degrading bacterium *Acinetobacter* sp. Tol 5, isolated from an exhaust gas biofiltration unit, is a promising chassis candidate for gas-phase bioprocesses (Hori et al., 2001). Tol 5 exhibits an auto-agglutination nature and high adhesiveness to various types of solid surfaces, mediated by the nanofiber protein AtaA on its cell surface (Ishikawa et al., 2012). This adhesiveness allows Tol 5 to be immobilized on carriers without chemical treatment, enabling its use as a whole-cell biocatalyst in bioprocesses (Yoshimoto et al., 2017). Utilizing this unique property, we previously developed a gas-phase bioprocess in which Tol 5 cells immobilized on the surface of a porous carrier were supplied with volatilized substrates in the absence of bulk water (Usami et al., 2020). This approach enabled efficient conversion of poorly water-soluble geraniol into (*E*)-geranic acid, a high-value monoterpenoid. Furthermore, recent genomic and transcriptomic analyses of Tol 5 have revealed its diverse catabolic pathways for volatile compounds and provided a fundamental basis for metabolic engineering (Inoue et al., 2026).

Microorganisms in gaseous environments experience physicochemical conditions different from those in aqueous media because of the absence of bulk water. Survival and metabolic activity of microorganisms under severely limited water availability have been extensively investigated in natural habitats such as soil, phyllosphere, lithic surfaces, and aerosols, as well as in engineered environments such as biofiltration units (Chan et al., 2012; Tecon and Or, 2017; Bryan et al., 2019; Flemming and Wuertz, 2019; Grinberg et al., 2019). In clinical settings, microbial survival strategies on dry surfaces have also been studied in the context of preventing pathogen transmission (Jawad et al., 1998; Lucidi et al., 2025). Bacterial cells in these environments experience water stress due to reduced water activity, as well as osmotic stress arising from increased solute concentrations. In addition, the absence of bulk water interrupts the continuous supply of exogenous nutrients, which may render intracellular reserves important for sustaining metabolism. Moreover, the higher oxygen availability, particularly in aerobic gas-phase bioprocesses, increases the potential for reactive oxygen species (ROS) generation, imposing oxidative stress on cells (Imlay, 2013). Bacteria exhibit various responses to counteract these environmental stresses, including synthesis of compatible solutes and antioxidants (Brown et al., 2003; Romantsov et al., 2009), modulation of cell membrane composition (Brown et al., 2003; Romantsov et al., 2009), biofilm formation (Flemming and Wuertz, 2019), and transition into a dormant state (Lebre et al., 2017; Pazos-Rojas et al., 2019; Lucidi et al., 2025). Such physiological adaptations are inevitably accompanied by alterations in cellular metabolism. However, while previous studies have primarily focused on survival and dormancy of pathogens, the metabolic adaptations that sustain catalytic activity in the gas phase remain poorly understood.

In this study, we investigated the metabolic alterations in Tol 5 cells degrading toluene in the aqueous and gas phases. Exploiting the adhesiveness of this strain, we assigned cell-immobilized carriers from the same cultures to the gas phase or to nitrogen-free aqueous media, so that the conditions differed only in phase and nutrient availability (Fig. 1A). Through an integrated analysis of metabolomics, lipidomics, and transcriptomics data, we characterized the metabolic features associated with the gas-phase condition. Elucidating these features provides a basis for the rational design of gas-phase bioprocesses.

**Figure 1.**
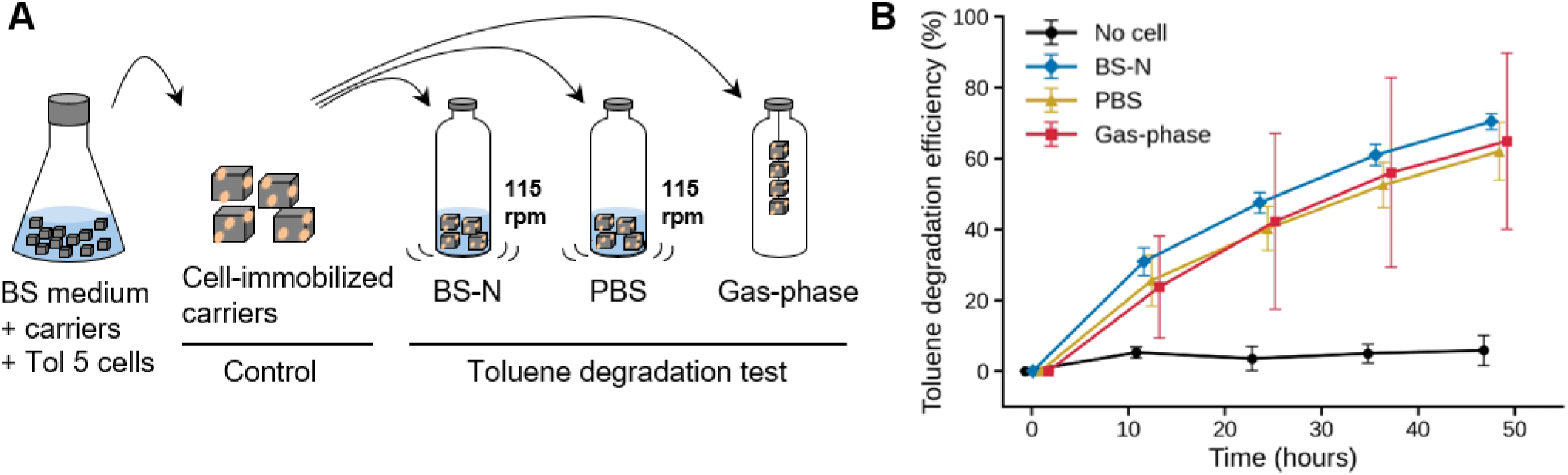
Experimental setup and time course of toluene degradation. (A) Experimental setup for the toluene degradation assay in the aqueous- and gas-phase conditions. (B) Toluene degradation assay. Toluene degradation was monitored by measuring the decrease in headspace toluene concentration after sealing cell-immobilized carriers in vials. The y-axis represents the toluene degradation efficiency, calculated relative to the headspace concentration measured at 30 min after toluene addition. The x-axis represents the incubation time. Data are presented as mean ± SD (n = 6 biological replicates for inoculated conditions; n = 3 for cell-free control).

## 2 Materials and methods

### 2.1 Culture conditions

*Acinetobacter* sp. Tol 5 was precultured overnight in a 50-mL centrifuge tube containing 4 mL of Lysogeny broth (LB) medium supplemented with 2 µL of toluene. The cells were washed with an equal volume of basal salt (BS) medium (Hori et al., 2001) and then inoculated (1:100) into a 500-mL baffled flask containing 100 mL of BS medium supplemented with 50 µL of toluene and 800 µL of 50% sodium DL-lactate. Sixteen cubic pieces of polyurethane (PU) foam (10 × 10 × 10 mm, CFH-20, 20 pores/25 mm; Inoac Corporation, Nagoya, Japan) were added as carriers for cell immobilization. The flask was capped with a Viton stopper and incubated at 28 °C with shaking at 115 rpm for 24 h. After cultivation, the PU carriers were retrieved, gently pressed against filter papers to remove excess liquid, and then subjected to various assays.

For microscopy, PU carriers bearing immobilized cells were sliced into 1-mm-thick sections and immersed in 4% paraformaldehyde phosphate buffer solution (FUJIFILM Wako, Osaka, Japan) for 15 min. After the sections were washed twice with phosphate-buffered saline (PBS; Nippon Gene, Tokyo, Japan), the cells on the sections were stained with 50 µM propidium iodide (Invitrogen, Waltham, MA, USA) in 50 mM HEPES buffer (pH 7.2) for 15 min. The sections were washed with PBS and observed under a digital microscope (KEYENCE, Osaka, Japan).

For colony formation assay, carriers bearing immobilized cells were collected after cultivation or after 48 h of the toluene degradation assay. Cells were detached from the carriers by shaking in 20 mL of 1% Casamino Acids technical grade (CA-T; Becton, Dickinson and Company, Franklin Lakes, NJ, USA) solution, which disperses AtaA-mediated cell aggregates (Ohara et al., 2019). Serial dilutions of the cell suspension were spotted onto LB agar plates and incubated at 28 °C overnight.

### 2.2 Toluene degradation assay

Bacterial cells were grown and immobilized on PU carriers in six independent cultures. The carriers from each culture were then divided among the aqueous- and gas-phase conditions, with four carriers assigned to each condition. For the aqueous-phase conditions, four PU carriers were immersed in each 100-mL vial containing 20 mL of modified BS medium lacking a nitrogen source (BS-N; BS containing 2.7 g L^−1^ K_2_SO_4_ instead of [NH_4_]_2_SO_4_) (Watanabe et al., 2008) or PBS. For the gas-phase condition, four PU carriers were suspended in the headspace of the vials using a stainless steel wire, as described previously (Usami et al., 2020). The vials were sealed with Teflon-laminated caps (5-112-7; Maruemu, Osaka, Japan), and 10 µL of toluene was added to the vial headspace using a gas-tight syringe (5181-8809; Agilent, Santa Clara, CA, USA). For the aqueous-phase conditions, the vials were incubated at 28 °C with shaking at 115 rpm. For the gas-phase condition, the vials were incubated at 28 °C without shaking.

Toluene in the headspace was quantified using a gas chromatograph equipped with a flame ionization detector (GC-FID) (GC-2014; Shimadzu, Kyoto, Japan). An HP-5ms Ultra Inert Column (30 m length, 0.25 mm inner diameter; Agilent, Santa Clara, CA, USA) was used with nitrogen as the carrier gas. A 50-µL aliquot of headspace gas was sampled from the vial using a gas-tight syringe (1710N PST-2; Hamilton, Reno, NV, USA) and injected into the GC-FID system. The oven temperature was maintained at 50 °C for 1 min and then increased to 80 °C at a rate of 20 °C min^−1^. The split ratio was 5:1, and the total gas flow rate was set to 19 mL min^−1^.

### 2.3 Sample preparation for LC–MS/MS analysis

Samples for the omics analyses were prepared from three independent cultures. After cultivation or after 24 h of the toluene degradation assay, cells were detached from the PU carriers by vigorous shaking in 5 mL of deionized water, in which cells immobilized via AtaA detach from the carriers and disperse (Yoshimoto et al., 2017). The cells were then harvested by centrifugation (10,000 × *g*, 4 °C, 1 min) and stored at −80 °C. Cells collected immediately after cultivation served as the control. Metabolite extraction from bacterial cell pellets was performed on ice using the Bligh and Dyer method (Bligh and Dyer, 1959). Dry cell pellets (5 mg) were mixed with 1,000 µL of ice-cold extraction solvent (methanol:chloroform:water = 10:4:4, v/v/v), vortexed for 1 min, and sonicated for 10 min. The samples were centrifuged at 16,000 × *g* for 5 min at 4 °C, and 900 µL of the resulting supernatant was transferred to a new tube. Phase separation was induced by adding 300 µL of chloroform and 252 µL of water, followed by vigorous shaking for 10 s. After centrifugation at 16,000 × *g* for 3 min at 4 °C, the upper aqueous phase (hydrophilic metabolites) and the lower organic phase (lipid metabolites) were collected into separate tubes and dried using a SpeedVac concentrator for 2.5 h. The upper phase was used for hydrophilic metabolomics analysis. Dried aqueous extracts were reconstituted in 50 µL of 5% methanol containing 0.2% formic acid and a mixture of internal standards (IS) (see Table S1 for details). The lower phase (organic phase) was used for lipidome analysis. Dried organic extracts were reconstituted in methanol containing IS, including EquiSPLASH LIPIDOMIX (Avanti Polar Lipids, USA), free fatty acid 16:0-d_3_ and 18:0-d_3_ (SRL, Canada) (see Table S2 for details). Reconstituted samples were centrifuged at 16,000 × *g* for 3 min at 4 °C, and 45 µL of the supernatant was collected. Of this volume, 30 µL was used for sample analysis, and 10 µL was pooled to prepare quality-control (QC) samples. The final aliquots were transferred to LC/MS vials (Agilent Technologies, Santa Clara, California, USA) for analysis. Hydrophilic metabolites in biological samples were analyzed in MS1 scanning mode, while lipid measurements were performed using data-dependent acquisition (DDA). QC samples were analyzed using data-independent acquisition (DIA), as described below.

### 2.4 LC–MS/MS measurement for hydrophilic metabolomics

Hydrophilic metabolites were analyzed according to a previously reported method (Lopes et al., 2021). The LC separation was performed on a 1290 Infinity III LC system (Agilent Technologies, Santa Clara, California, USA). Metabolites were separated using an ACQUITY UPLC HSS T3 Column (2.1 mm × 50 mm, 1.8 µm) (Waters Corporation, Milford, Massachusetts, USA). Throughout the LC–MS measurement, the column temperature was maintained at 45 °C, and the flow rate was set to 0.3 mL min^−1^. The mobile phases comprised solvent (A) water with 0.2% formic acid and solvent (B) methanol with 0.1 % formic acid. A sample volume of 1 μL was used for the injection. The separation was performed under the following gradient: 0 min 0.1% (B), 0.5 min 0.1% (B), 2.0 min 60% (B), 2.3 min 95% (B), 3.0 min 95% (B), 3.1 min 0.1% (B), 4.5 min 0.1% (B), 4.6 min 0.1% (B), and 5.5 min 0.1% (B). The injection and LC gradients required 30 s and 5.5 min, respectively; therefore, the injection-to-injection time was 6.0 min. The temperature in the sample rack was maintained at 4 °C.

The MS detection was performed using QTOF-MS (6546 LC/Q-TOF system; Agilent Technologies, Santa Clara, California, USA). The DDA method for QC samples was used in both positive and negative ion modes with the following parameters: MS1 accumulation time, 200 ms; MS2 accumulation time, 100 ms; Q1 resolution, narrow (∼1.3 *m/z*); and no exclusion range. The following settings were used for positive ion mode: MS1 mass ranges, *m/z* 50–950; MS2 mass ranges, *m/z* 50– 950; reference masses, *m/z* 121.050873 and 922.009798; and exclusion lists, *m/z* 121.050873, *m/z* 149.02332, and *m/z* 922.009798. The following settings were used for negative ion mode: MS1 mass ranges, *m/z* 50–970; MS2 mass ranges, *m/z* 50–950; reference masses, *m/z* 119.03632 and 966.000725; and exclusion lists, *m/z* 119.03632 and *m/z* 966.000725. Ion source parameters for both positive and negative ion modes were as follows: ionization, ESI; source temperature, 225 °C; nebulizer gas, 50 psi; sheath gas, 12 L/min. Collision energies were set to 20 eV for positive ion mode and 30 eV for negative ion mode. MS1 scanning mode was set in both positive and negative ion modes for biological samples as follows: mass range, *m/z* 50–950; acquisition rate, 5 spectra/s; acquisition time, 200 ms; transients/spectrum, 1800. The other settings were the same as those used for DDA, except for the MS/MS acquisition parameters. The DIA method was set in both positive and negative ion modes as follows: MS1 accumulation time, 200 ms; MS2 accumulation time, 100 ms; Q1 window, 46 Da; and MS1 mass range, *m/z* 50–950. The other settings were the same as those used for DDA. Five different DIA settings were prepared in which the precursor scanning ranges for MS/MS acquisition were set in both ion modes to 50–230, 230–410, 410–590, 590–770, and 770–950 Da.

Biological samples were measured in MS1 scanning mode. QC samples were measured using both DIA and DDA modes to ensure comprehensive, high-quality MS/MS coverage. For DDA measurements, an iterative MS/MS method was employed to minimize redundant precursor fragmentation across repeated injections.

### 2.5 LC–MS/MS measurement for lipidomics

Lipidomics was performed according to a previously reported method (Tokiyoshi et al., 2024) using the same instrument as for hydrophilic metabolomics. Lipids were separated using an Imtakt UK-C18 MF column (50 × 2.0 mm, 3 µm) (Imtakt, Kyoto, Japan) equipped with a matching Imtakt guard column (5 × 2.0 mm) (Imtakt, Kyoto, Japan). Throughout the LC–MS measurement, the column temperature was maintained at 65 °C, and the flow rate was set to 0.6 mL min^−1^. The mobile phases comprised solvent (A) 1:1:3 (v/v/v) acetonitrile: methanol: water with ammonium acetate (5 mM) and ammonium fluoride (0.2 mM) and solvent (B) 1:9 (v/v) acetonitrile: isopropyl alcohol with ammonium acetate (5 mM) and ammonium fluoride (0.2 mM). A sample volume of 1 μL was used for the injection. The separation was performed under the following gradient: 0 min 0.1 % (B), 0.1 min 0.1 % (B), 0.2 min 15 % (B), 1.1 min 30 % (B), 1.4 min 48 % (B), 5.6 min 82 % (B), 6.9 min 99.9 % (B), 7.1 min 99.9 % (B), 7.2 min 0.1 % (B), and 8.6 min 0.1 % (B). The injection and LC gradients required 30 s and 8.6 min, respectively; therefore, the injection-to-injection time was 9.1 min. The temperature of the rack was maintained at 4 °C.

The DDA method was used in both positive and negative ion modes with the following parameters: MS1 and MS2 accumulation time, 100 ms; Q1 resolution, narrow (∼1.3 *m/z*). The following settings were used for positive ion mode: MS1 mass ranges, *m/z* 75–1250; MS2 mass ranges, *m/z* 50–1250; reference masses, *m/z* 121.050873 and 922.009798; exclusion range, *m/z* 75–300; and exclusion lists, *m/z* 922.009798. The following settings were used for negative ion mode: MS1 and MS2 mass ranges, *m/z* 75–1250; reference masses, *m/z* 119.03632 and 966.000725; exclusion range, *m/z* 75– 200; and exclusion lists, *m/z* 119.03632 and *m/z* 966.000725. Ion source parameters for positive/negative ion modes were as follows: ionization, ESI; source temperature, 250/300 °C; nebulizer gas, 50 psi (both modes); sheath gas, 12 L/min (both modes); collision energy, 20/30 eV. The DIA method was set in both positive and negative ion modes as follows: MS1 and MS2 accumulation time, 100 ms; and MS1 mass range, *m/z* 70–1250. Q1 windows were 63.6 Da (positive ion mode) and 70.5 Da (negative ion mode). The other settings were the same as those used for DDA. Five different DIA settings were prepared in which the precursor scanning ranges for MS/MS acquisition were set in positive ion mode to 300–490, 490–680, 680–870, 870–1060, and 1060–1250 Da. In negative ion mode, five different DIA settings were prepared in which the precursor scanning ranges for MS/MS acquisition were set to 200–410, 410–620, 620–830, 830–1040, and 1040–1250 Da. The DDA method was used to measure biological samples, and the DIA method was used to measure QC samples.

### 2.6 Data processing for metabolome and lipidome analyses

The mass spectrometry data were analyzed using MS-DIAL 5 (ver. 5.4.241021) (Takeda et al., 2024). MS-DIAL parameter settings were as follows: MS1 and MS2 data type, “centroid”; acquisition type, “DDA” and “SWATH”, according to the data type; max number of isotope recognitions, 2; the number of threads, 8; the minimum peak amplitude, 500; MS/MS absolute abundance cutoff, 0; MS/MS relative abundance cutoff, 0. The default parameter values were used for the others. Detailed parameter settings are provided in Supplementary Information. Lipid annotation was performed following a previously described protocol (Tsugawa et al., 2020). The predicted retention times in the lipidomics spectral database were adjusted to the LC–MS conditions used in this study by using the relationship between the retention time and the solvent B% values in the LC gradient condition. Hydrophilic metabolites were annotated with three MSP spectral libraries, following a previously reported strategy (Kiuchi et al., 2024). MSP1 contained metabolites with experimentally confirmed retention times (1311 records for positive ion mode, 833 records for negative ion mode). MSP2 contained metabolites for which retention times were predicted by a machine learning method (43870 records for positive ion mode, 26684 records for negative ion mode). MSP3 consisted of metabolites without retention time information (864 records for positive ion mode, 599 records for negative ion mode). All records contained MS/MS spectra. No normalization was applied to the hydrophilic metabolite data because the cell weight was identical across samples; instead, QC drift correction was performed using “notame” (Klåvus et al., 2020). For lipidomics data, normalization based on internal standards was performed using the EquiSPLASH lipidomix normalization function in MS-DIAL 5. Table S3 lists the pairs of internal standards and lipid subclasses used for normalization. Inter-sample correction was performed using the median correction method. Source code is provided as a Source Data file. Metabolomics data were further classified according to annotation confidence levels based on the criteria proposed by Juliane Hollender (Schymanski et al., 2014). Only metabolites assigned to annotation levels 1 and 2 were used for downstream analyses, including multivariate analysis and pathway analysis.

### 2.7 Multivariate analysis for lipidome and metabolome

Principal component analysis (PCA) was performed to evaluate overall differences between groups. Prior to analysis, QC samples were excluded. PCA was conducted using the prcomp function in the R language (version 4.5.2). Hierarchical clustering analysis (HCA) was performed on metabolome and lipidome data after Pareto scaling to reduce the influence of large variance while preserving biologically relevant differences. Clustering was performed using Euclidean distance and Ward’s minimum variance method (Ward.D2). Sample class annotations were incorporated into the heatmaps for visualization. Metabolome data were partitioned into six clusters, while lipidome data were partitioned into five clusters based on the hierarchical tree structure. All visualizations, including heatmaps and dendrograms, were generated using the “pheatmap”, “ggplot2”, and “ggdendro” packages in R (version 4.5.2). Enrichment analysis of lipidome was performed using LION (https://www.lipidontology.com/index.html) (Molenaar et al., 2019), where the ranking mode was utilized. For local statistics to rank input identifiers, “2-LOG[fold change] (2 conditions)” was selected. Enrichment analysis of metabolomics was performed using MetaboAnalyst 6.0 (https://www.metaboanalyst.ca/home.xhtml) (Pang et al., 2024), where the enrichment analysis mode was used. The following normalization preprocessing options were applied: sample normalization, none; data transformation, log2 transformation; data scaling, auto scaling. Kyoto Encyclopedia of Genes and Genomes (KEGG) database was selected as the reference pathway. Visualization of individual metabolite levels for pathway mapping was performed using the “ggplot2” package in R (version 4.5.3). Statistical comparisons against the control group were conducted using Dunnett’s test, implemented via the “multcomp” package in R (version 4.5.3).

### 2.8 RNA sequencing and data processing for transcriptome analysis

Total RNA was extracted using the Cica geneus RNA Prep Kit (for Tissue) (Kanto Chemical, Tokyo, Japan) following the manufacturer’s protocol. Ribosomal RNA (rRNA) was depleted using the NEBNext rRNA Depletion Kit (Bacteria) (New England Biolabs, Ipswich, MA, USA), and cDNA libraries were prepared with the NEBNext Ultra II RNA Library Prep Kit for Illumina (New England Biolabs, Ipswich, MA, USA). Sequencing was performed on an Illumina NextSeq 550 system (single-end 81-bp reads). Raw reads were quality-filtered using fastp (version 0.23.2) (Chen et al., 2018) and mapped to the Tol 5 genome (accession numbers: AP024708 and AP024709) using Bowtie2 (version 2.5.1) (Langmead and Salzberg, 2012) with default parameters. Mapped reads were counted using featureCounts (version 2.0.6) (Liao et al., 2014). Statistical analysis was performed using the “edgeR” package in the R language (version 4.4.1). Genes with extremely low counts were filtered out using the filterByExpr function. Normalization was performed using the trimmed mean of M-values (TMM) method implemented in the calcNormFactors function.

### 2.9 Transcriptomic data analysis

PCA was conducted on log_2_ counts per million (CPM) using the prcomp function in the R language. Differential gene expression analysis was conducted using the quasi-likelihood pipeline of the “edgeR” package. Dispersion estimates were obtained using the estimateDisp function, and model fitting was performed with the glmQLFit function. Differential gene expression was assessed using the glmQLFTest function, comparing the BS-N, PBS, and gas-phase conditions with the control condition. The full results are provided in Table S4.

Over-representation analysis (ORA) was performed using the “clusterProfiler” package in the R language (Yu et al., 2012). Differentially expressed genes identified from the gas-phase vs. BS-N and gas-phase vs. PBS comparisons (|log_2_ fold change [FC]| > 1, false discovery rate [FDR] < 0.01, log_2_ CPM > 3) were analyzed separately for up- and downregulated genes against all detected genes of Tol 5. Enrichment was assessed for KEGG Orthology pathways and Clusters of Orthologous Genes (COG) categories, using a minimum gene set size of 5 (KEGG) or 2 (COG), a maximum gene set size of 5000, and Benjamini–Hochberg *p*-value adjustment. The results were visualized using the “ggplot2” package in the R language.

## 3 Results

### 3.1 Toluene degradation capability of Tol 5 in the aqueous and gas phases

To investigate the metabolic differences between metabolically active Tol 5 cells in conventional aqueous-phase bioprocesses and those in gas-phase bioprocesses without a surrounding bulk aqueous medium, we first confirmed their ability to metabolize toluene, a carbon source that can be supplied in both phases. Tol 5 cells were initially cultured in BS medium containing the PU carriers, with toluene supplied as a carbon source. Due to the high adhesiveness of the cells, they adhered to the carriers during growth, allowing the cell-immobilized carriers from the same cultures to be assigned to the aqueous- or gas-phase conditions without cell detachment (Figs. 1 and S1A). Microscopic observation showed that the cells were distributed over the foam surface, while the open pore structure of the carrier left internal voids through which liquid or gas could pass (Figs. S1B and C). To exclude the effects of cell growth, the aqueous-phase conditions comprised two nitrogen-free solutions: BS-N, a modified BS medium lacking a nitrogen source, and PBS. The gas-phase process in low-water environments was established by suspending the carriers in the headspace of a vial, following the previous study (Usami et al., 2020). In the gas-phase condition, each set of four carriers retained 5.8 ± 1.7 mg of dry cells and 378.2 ± 60.2 mg of water (mean ± standard deviation, n = 6 biological replicates) after removal of excess liquid (Figs. S1C and D). The water content of the cell-immobilized carriers remained essentially unchanged throughout the reaction (Fig. S2).

Ten microliters of toluene was added to the vial headspace in each condition, where it evaporated immediately, and its consumption was monitored. Based on the dimensionless Henry’s constant of toluene in BS-N medium (0.315 at 28 °C) (Morono et al., 2004), which was also applied to PBS because of its comparable ionic strength, 44% of the added toluene was estimated to dissolve in the aqueous phase in the aqueous conditions, giving a calculated concentration of 191 mg L^−^¹ (2.1 mM). In the gas-phase condition, 98.8% of the substrate resided in the gas phase at 90 mg L^−^¹, with the remaining 1.2% dissolved in the small amount of water retained by the carriers, corresponding to an aqueous concentration of 285 mg L^−^¹ (3.1 mM). In all conditions, toluene in the headspace decreased progressively over 48 h, confirming that the cells were metabolically active in both the aqueous and gas phases; toluene consumption continued beyond this time point in all conditions (Fig. 1B). At 24 h, the degradation efficiency was 47.5 ± 3.0%, 40.2 ± 6.2%, and 42.3 ± 24.8% in BS-N, PBS, and the gas phase, respectively, indicating that degradation was still in progress at this time point in every condition. Cell viability, assessed by colony formation at 48 h, was comparable among the conditions (Fig. S3). We therefore focused on the cellular state at 24 h, when toluene degradation was actively proceeding in all conditions, and the collected cells were subjected to metabolome, lipidome, and transcriptome analyses.

### 3.2 Metabolomic alterations in Tol 5 in the aqueous and gas phases

The hydrophilic metabolites were extracted and quantified by LC–MS/MS, yielding 97 metabolites detected across all conditions. PCA of the detected metabolites clearly separated cells collected immediately after cultivation in BS medium (control) from cells in the three conditions during the toluene degradation assay (BS-N, PBS, and gas phase) (Fig. 2A). Specifically, PC1, which accounted for 38.5% of the variance, separated the control samples from the others, consistent with a major metabolic alteration accompanying the transition from growth on toluene in nitrogen-containing BS medium to toluene degradation under nitrogen-free resting cell conditions. When PCA was repeated using only samples collected under the three toluene degradation conditions, PC3 separated the aqueous-phase samples (BS-N and PBS) from the gas-phase samples (Fig. 2B), with the gas-phase samples on the negative side of PC3. This separation was reproducible across biological replicates despite variation in toluene degradation efficiency, indicating a gas-phase-specific metabolic state. Metabolites with large negative PC3 loadings, associated with higher accumulation levels in the gas phase, included key intermediates of glycolysis/gluconeogenesis and the tricarboxylic acid (TCA) cycle, such as 3-phosphoglyceric acid, phosphoenolpyruvic acid, and isocitric acid (Fig. 2C). Intermediates of nucleic acid metabolism, including cytidine 5’-monophosphate (CMP), uridine 5’-monophosphate (UMP), and guanosine 5’-monophosphate (GMP), and of amino acid metabolism such as glutamic acid and argininosuccinic acid, also showed large negative loadings. HCA reproduced this separation and resolved six clusters (Fig. S4). Cluster 5 was specifically reduced under the gas phase condition and comprised di- and tripeptides and pyridine derivatives, matching the gas-phase-specific reductions inferred from the PC3 loadings. The gas-phase cells grouped most closely with those in PBS, the most nutrient-limited aqueous condition.

**Figure 2.**
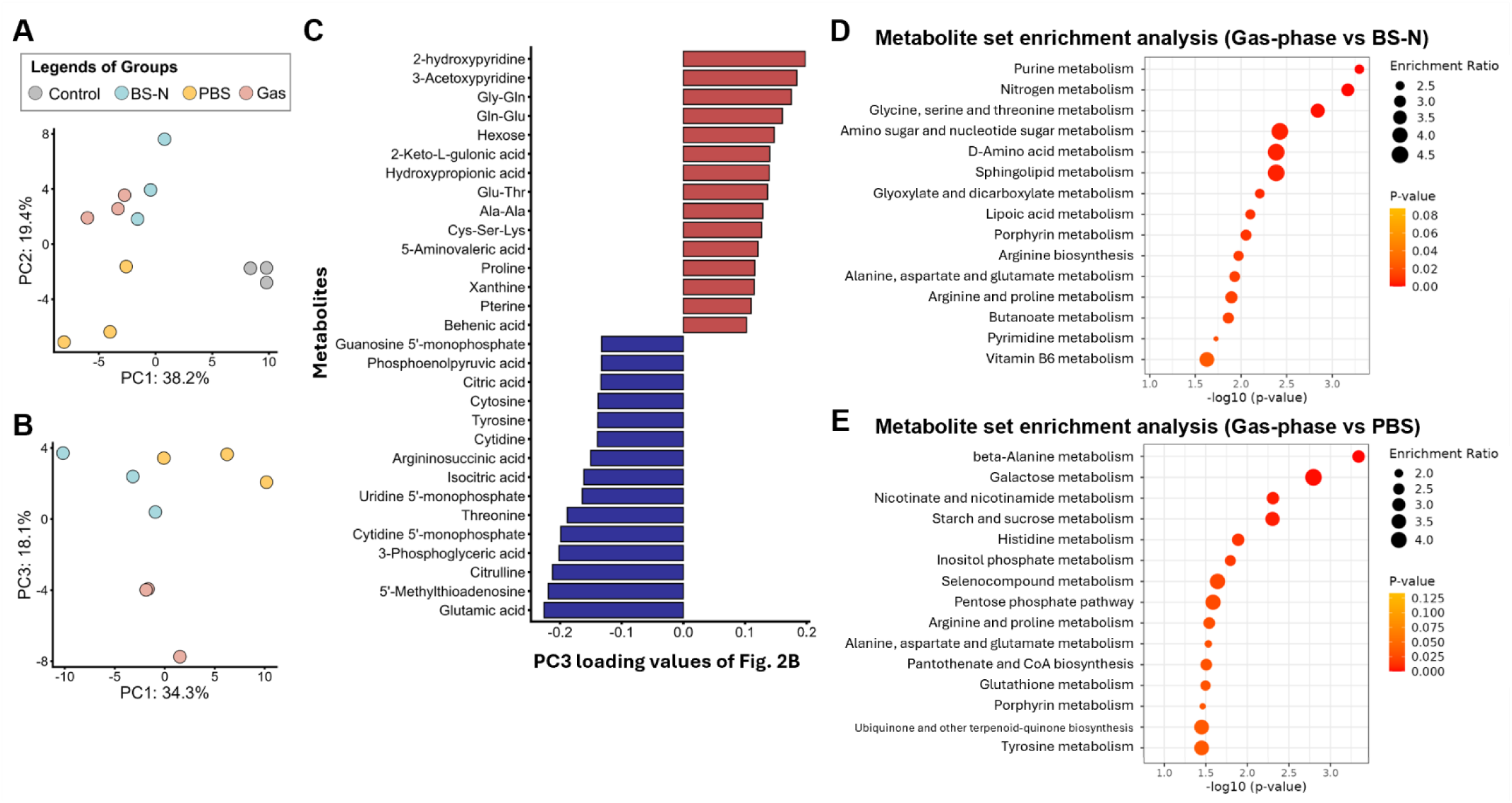
Metabolome data mining for Tol 5 cells in the aqueous- and gas-phase conditions. (A) PCA score plot based on the metabolome profiles (n = 3 biological replicates). (B) PCA score plot with the control samples excluded. (C) Loading values of the metabolites contributing most strongly to PC3 shown in (B). The bar plot displays the top 15 positive (red) and top 15 negative (blue) loadings. The y-axis represents the individual metabolites, and the x-axis indicates the loading value on PC3. (D, E) Enrichment analysis performed using MetaboAnalyst 6.0, showing the top 15 enriched pathways for (D) the gas phase compared with BS-N and for (E) the gas phase compared with PBS.

To identify metabolic processes specifically altered under the gas-phase condition, we performed an enrichment analysis comparing cells in this condition with those in BS-N and PBS (Figs. 2D and E). While multiple metabolic processes were enriched, various amino acid metabolic pathways consistently ranked at the top in both comparisons. In addition, metabolic processes related to nucleic acids and cofactors were identified. Most of these metabolic pathways involve nitrogen-containing metabolites. Notably, none of the three conditions contained an external nitrogen source, and the pathways differing between conditions therefore operated on intracellular nitrogen.

### 3.3 Lipidomic alterations in Tol 5 in the aqueous and gas phases

Lipid components were extracted and quantified by LC–MS/MS, yielding 108 lipid species detected across all conditions. Phosphatidylglycerol (PG) and phosphatidylethanolamine (PE) were the major phospholipids in Tol 5, consisting mainly of species with acyl chain lengths of 16 to 18 carbons and a degree of unsaturation of 0 to 1 (16:0, 16:1, 18:0, and 18:1) (Fig. S5). This lipid profile resembles that of typical Gram-negative bacteria (Walczak-Skierska et al., 2024). We also detected lysophosphatidylethanolamine (LPE), cardiolipin (CL), monolysocardiolipin (MLCL), and a triacylated glycerophosphoglycerol species annotated as hemibis(monoacylglycero)phosphate (HBMP), which is a structural isomer of acylphosphatidylglycerol (acyl-PG) and could not be distinguished from it under the present LC–MS/MS conditions. Phosphatidic acid (PA), the common intermediate of glycerophospholipid synthesis, was also detected. Among neutral lipids, monoacylglycerol (MG), diacylglycerol (DG), triacylglycerol (TG), and wax esters (WE) were detected.

PCA of the lipid abundance profiles separated samples collected under the three toluene degradation conditions (BS-N, PBS, and gas phase) from the control samples (Fig. 3A). When PCA was repeated using only samples collected under the three toluene degradation conditions, PC1 separated the gas-phase samples from those in PBS and BS-N, with the gas-phase samples on the negative side of the axis (Fig. 3B). Lipid species with large negative PC1 loadings, associated with higher accumulation levels in the gas phase, were dominated by PE, PG, and several coenzyme Q (CoQ) species, whereas those with large positive loadings, associated with lower accumulation levels in the gas phase, were dominated by neutral lipids, particularly diverse DG and TG species. HCA reproduced this pattern and resolved five clusters, of which Clusters 3 and 5 were specific to the gas-phase condition (Fig. S6). Cluster 3, comprising species at higher levels in the gas phase, contained PG, PE, and CoQ, together with several LPE and HBMP species. Cluster 5, comprising species at lower levels in the gas phase, contained TGs and WEs across all chain lengths; MLCL followed the same pattern. In the HCA, the gas-phase cells grouped most closely with those in BS-N rather than PBS, in contrast to the metabolome, in which the gas-phase cells grouped with those in PBS.

**Figure 3.**
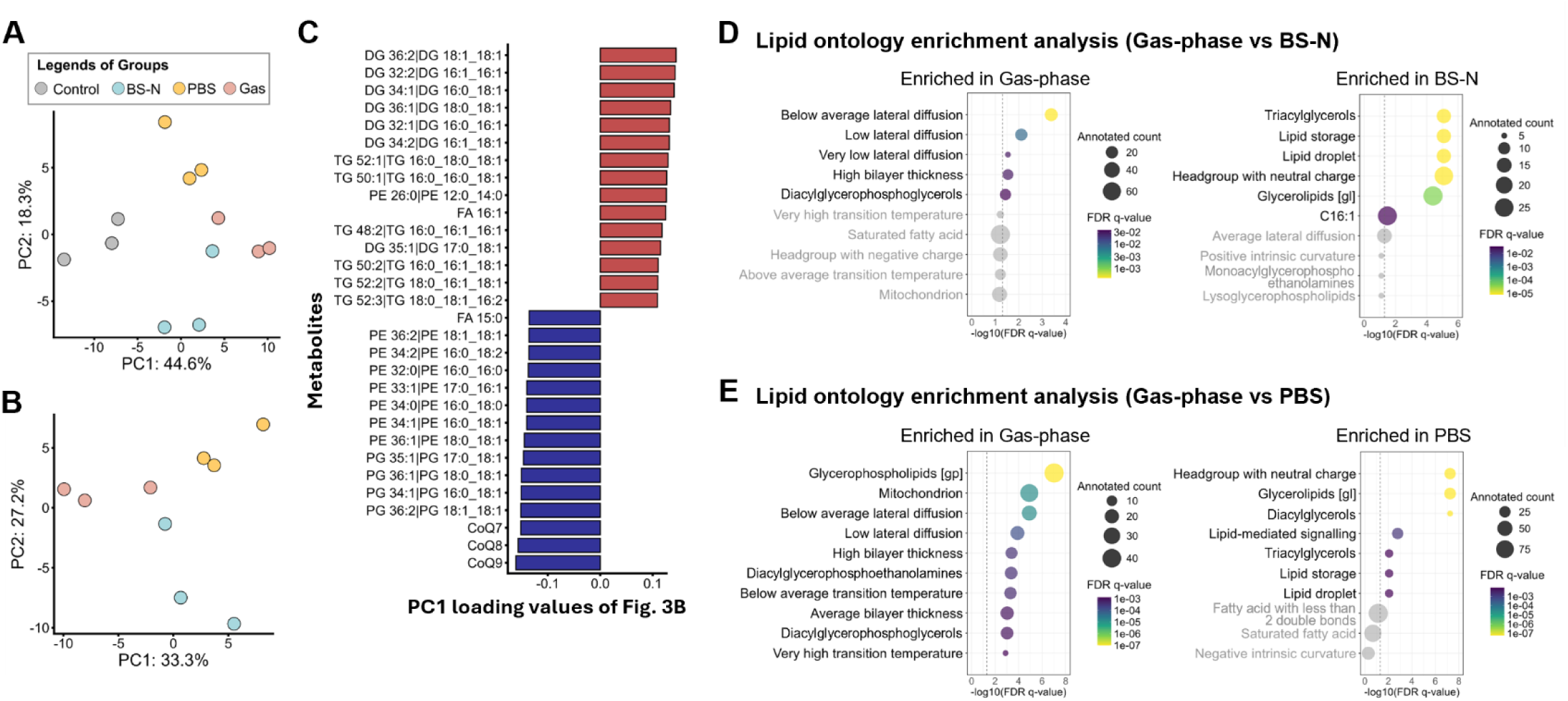
Lipidome data mining for Tol 5 cells in the aqueous- and gas-phase conditions. (A) PCA score plot based on the lipidome profiles (n = 3 biological replicates). (B) PCA score plot with the control samples excluded. (C) Loading values of the lipid species contributing most strongly to PC1 in the PCA of the three toluene degradation conditions shown in (B). The bar plot displays the top 15 positive (red) and top 15 negative (blue) loadings. The y-axis lists the individual lipid species, and the x-axis indicates the loading value on PC1. (D, E) Lipid ontology enrichment analysis, showing the top 10 enriched LION terms for (D) the gas phase compared with BS-N and for (E) the gas phase compared with PBS. LION terms without statistically significant enrichment (FDR > 0.05) are shown as gray bubbles, and the FDR threshold of 0.05 is indicated by a gray dashed vertical line.

Lipid ontology enrichment analysis comparing cells under the gas-phase condition with those in BS-N and PBS identified alterations in membrane and storage lipids (Figs. 3D and E). In both comparisons, lipid species enriched under the gas-phase condition were associated with lower lateral diffusion and increased bilayer thickness, consistent with the higher PE and PG levels and the shifts in their acyl chain lengths (Fig. S5). Conversely, the depleted species were mainly storage lipids, specifically TGs and DGs (Figs. 3D and E). Collectively, the lipidome data indicate remodeling of membrane lipids and a reduction in storage lipids specific to the gas-phase condition.

### 3.4 Transcriptomic alterations in Tol 5 in the aqueous and gas phases

To elucidate the gene expression profiles associated with the observed metabolomic and lipidomic alterations, and to investigate cellular responses not detectable by these analyses, transcriptome analysis was performed on cells under the same experimental conditions. Sequencing reads were mapped to the Tol 5 reference genome, and the expression levels of 3,899 genes were quantified. PCA of gene expression separated samples collected under the three toluene degradation conditions (BS-N, PBS, and gas phase) from the control samples, similar to the metabolome and lipidome data (Fig. 4A). When the analysis was restricted to the three toluene degradation conditions, PC1 separated the three conditions from one another, with the gas-phase cells on the positive side of the axis (Fig. 4B). Among the COG categories, genes with large positive PC1 loadings, associated with relatively higher expression in the gas phase, were enriched in lipid and amino acid metabolism, whereas those with large negative loadings, associated with relatively lower expression in the gas phase, were enriched in cellular processes and information storage (Figs. S7A–C).

**Figure 4.**
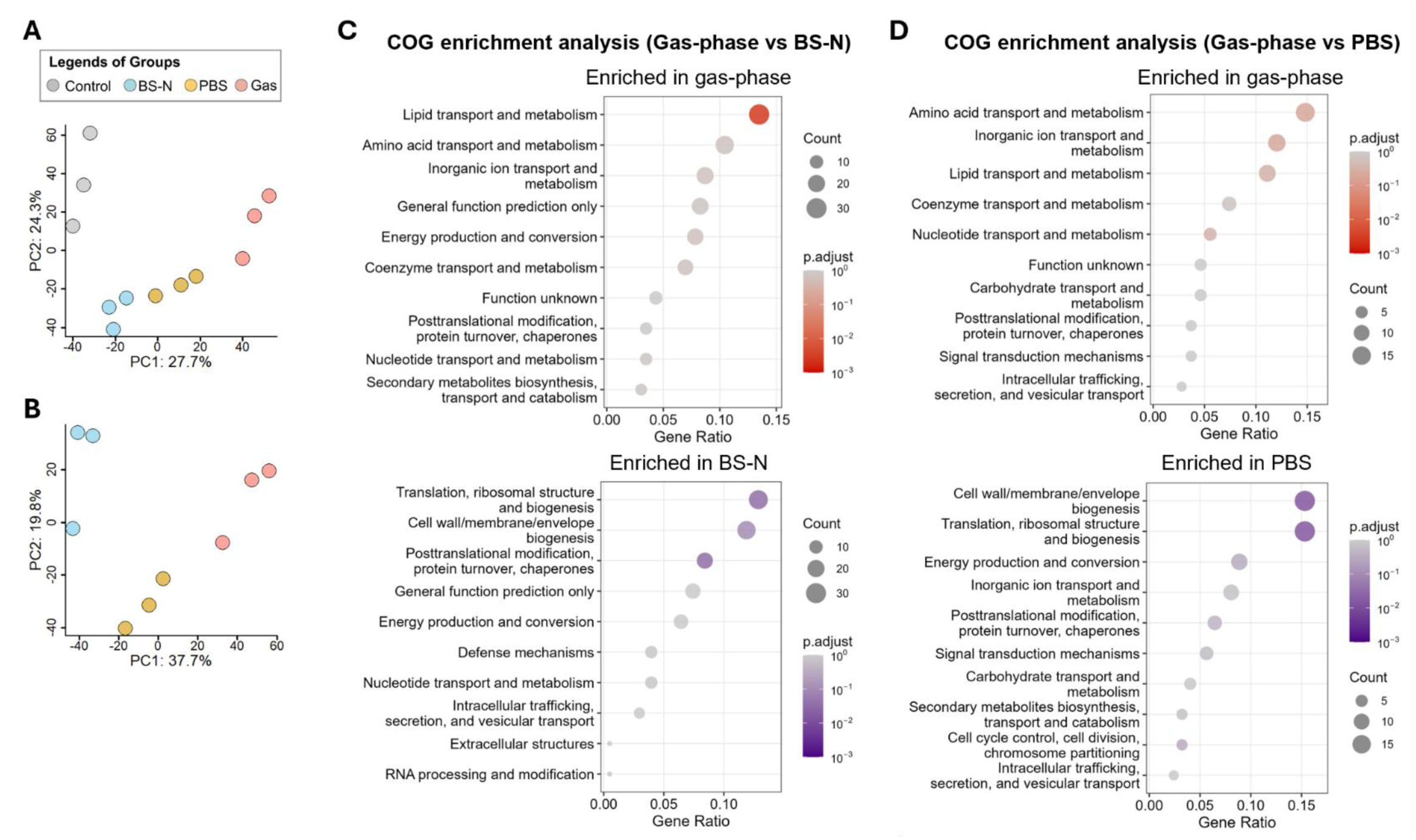
Transcriptome data mining for Tol 5 cells in the aqueous- and gas-phase conditions. (A) PCA score plot based on the transcriptome profiles (n = 3 biological replicates). (B) PCA score plot with the control samples excluded. (C, D) ORA of differentially expressed genes across COG functional categories, performed using clusterProfiler for (C) the gas phase compared with BS-N and (D) the gas phase compared with PBS. In each panel, categories enriched among genes upregulated in the gas phase (upper) and among genes upregulated in the corresponding aqueous condition, i.e., downregulated in the gas phase (lower), are shown separately. The x-axis indicates the gene ratio, the dot size represents the number of differentially expressed genes assigned to each category, and the color gradient indicates the Benjamini–Hochberg adjusted *p* value.

Differentially expressed genes were defined as those with |log_2_ FC| > 1, FDR < 0.01, and log_2_ CPM > 3. ORA of differentially expressed genes across COG functional categories was performed to compare the gas-phase condition with BS-N and PBS. In both comparisons, lipid transport and metabolism and amino acid transport and metabolism were the most highly enriched categories among genes upregulated in the gas phase, together with inorganic ion, nucleotide, and coenzyme transport and metabolism (Figs. 4C and D). Conversely, categories associated with translation and ribosomal biogenesis and with cell wall/membrane/envelope biogenesis were most strongly enriched among genes downregulated in the gas-phase condition. The concurrent enrichment of the amino acid and nucleotide categories suggests a shift in nitrogen-source utilization under the gas-phase condition, consistent with the nitrogen-related alterations inferred from the metabolome analysis (Figs. 2D and E). In comparison with both BS-N and PBS, KEGG pathway ORA also highlighted several amino acid metabolic pathways, including phenylalanine metabolism and valine, leucine, and isoleucine degradation, among genes upregulated in the gas phase (Fig. S8). The enrichment of lipid transport and metabolism likewise corresponds to the storage lipid changes observed in the lipidome (Figs. 3C–E).

### 3.5 Maintenance of the glutamate pool and mobilization of endogenous nitrogen in the gas phase

Both the metabolome and transcriptome pointed to a reorganization of nitrogen and amino acid metabolism as a defining feature of the gas-phase condition. In particular, glutamate (glutamic acid) was a key metabolite characterizing the gas-phase-specific alteration (Fig. 2C). Because glutamate is the primary ammonia-assimilation product and the central amino-group donor in bacterial nitrogen metabolism (Schulz-Mirbach et al., 2022), we mapped the detected metabolites and gene expression levels onto the metabolic pathways of Tol 5 (Fig. 5).

**Figure 5.**
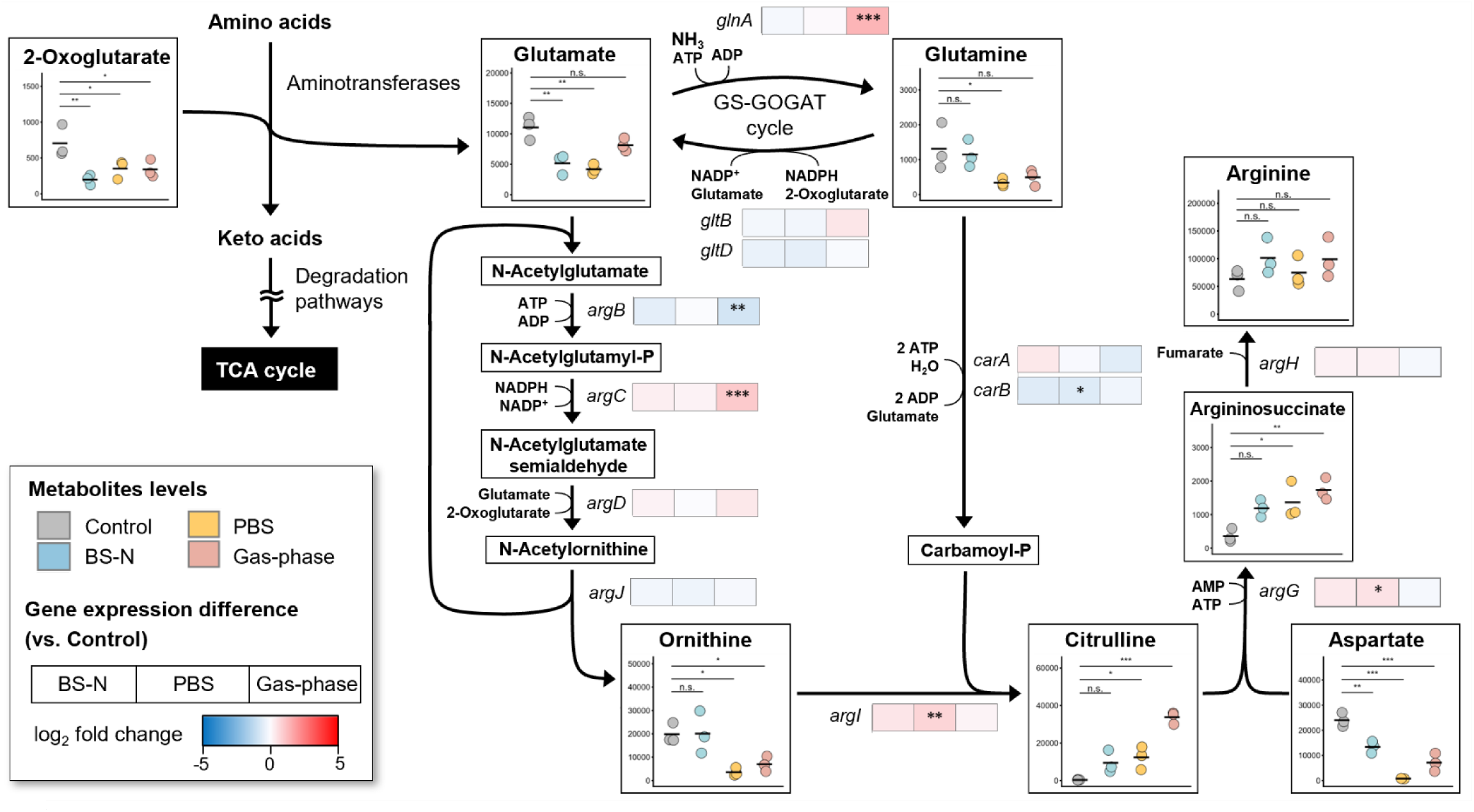
Metabolite levels and differential gene expression for glutamate metabolism. For metabolites, each dot represents the normalized peak height of one of three biological replicates from the control, BS-N, PBS, and gas-phase conditions, and the bars indicate the mean values. Asterisks indicate statistical significance based on adjusted *p* values obtained by Dunnett’s test (* *p* < 0.05, ** *p* < 0.01, *** *p* < 0.001, **** *p* < 0.0001), with the control condition as the reference. For gene expression, each box represents a gene, and the color gradient indicates the log_2_ FC of gene expression in the BS-N, PBS, and gas-phase conditions relative to the control condition. Asterisks within the boxes indicate statistical significance based on the FDR calculated in the differential gene expression analysis (* FDR < 0.05, ** FDR < 0.01, *** FDR < 0.001).

The glutamate pool was high in the control cells grown in nitrogen-replete BS medium and declined in the nitrogen-free aqueous conditions (BS-N and PBS). Despite the absence of external nitrogen sources, the glutamate pool in the gas-phase condition remained higher than those in BS-N and PBS. Among the detected amino acids, only glutamate was higher in the gas phase than in BS-N and PBS (Fig. S9). In the transcriptome, glutamine synthetase (GS, encoded by *glnA*), which together with glutamate synthase (GOGAT, encoded by *gltBD*) constitutes the GS–GOGAT ammonia-assimilation cycle, was strongly upregulated only in the gas phase.

Amino acid degradation pathways were also altered in the gas phase. Genes for the phenylacetic acid pathway (*paa* genes) and homogentisate pathway (*hmg* genes), which degrade the carbon skeletons of phenylalanine and tyrosine after their amino groups are transferred to 2-oxoglutarate, were prominently upregulated in the gas phase (Table S5). Because this transamination yields glutamate, the degradation of these aromatic amino acids represents a potential route for glutamate formation in the gas phase. The aminotransferase genes *ilvE*, *tyrB*, and *ybdL*, which catalyze the reversible transfer of amino groups between amino acids and 2-oxoglutarate/glutamate, were unchanged across the three conditions, indicating that these transamination steps were not transcriptionally regulated under the conditions tested.

Genes mediating the release of nitrogen from endogenous compounds were also upregulated in the gas phase. Xanthine, the branch point between purine salvage and degradation to urate, was specifically decreased in the gas phase, consistent with a shift from salvage toward degradation (Fig. S10). Downstream, genes for urate degradation (*aba2*, *hiuH*, *uao*, *cda1*, and *alc*), which convert urate to urea, and for urease (*ure* genes), which hydrolyzes urea to ammonia and carbon dioxide, were upregulated in all three conditions and most markedly in the gas phase (Fig. S10). Among the pyrimidines, UMP and CMP increased relative to the control in all three conditions, and the uracil degradation genes (*rut* genes), which also release ammonia, were upregulated (Fig. S11). Genes encoding nitrate reductase, which reduce nitrate to ammonia, were also upregulated in the gas phase despite the absence of their substrate in all conditions (Table S5).

Further downstream, glutamine, ornithine, and aspartate clearly decreased in PBS and the gas phase, whereas citrulline accumulated under the three toluene degradation conditions and was substantially higher in the gas phase than in BS-N and PBS (Fig. 5). Among the genes for citrulline biosynthesis from glutamate, N-acetyl-γ-glutamyl phosphate reductase (*argC*) was upregulated and N-acetylglutamate kinase (*argB*) was downregulated specifically in the gas phase. Argininosuccinate, synthesized from citrulline and aspartate, also increased under the three toluene degradation conditions, whereas the final product arginine did not differ across conditions.

### 3.6 Enhanced degradation of storage lipids and alterations in cell membrane composition in the gas phase

The lipidome analysis had shown altered storage- and membrane-lipid composition specific to the gas phase (Fig. 3). To better understand these changes, we integrated the transcriptomic data and mapped them onto the lipid metabolic pathways (Fig. 6).

**Figure 6.**
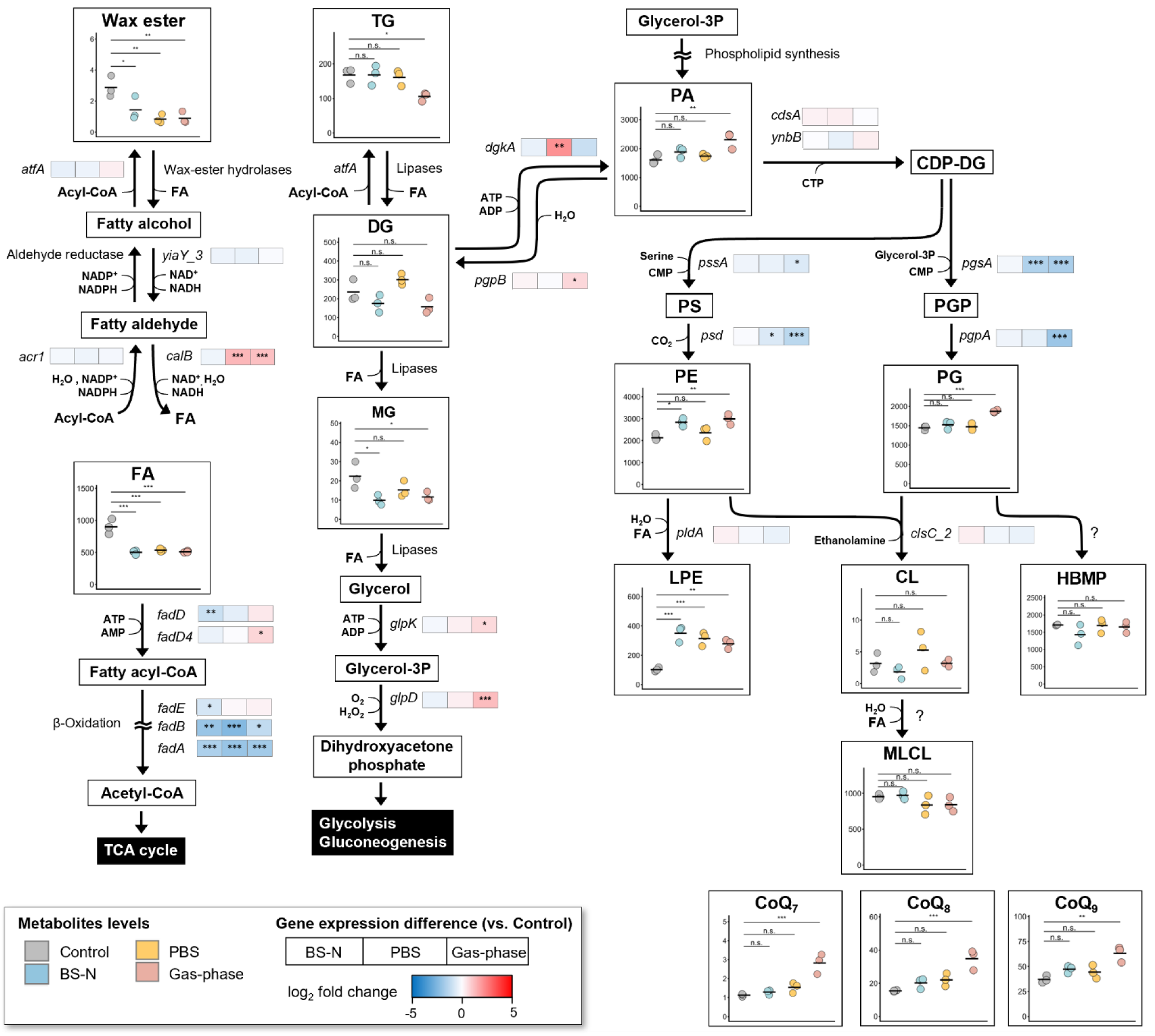
Metabolite levels and differential gene expression for lipid metabolism. For metabolites, each dot represents the normalized peak height of one of three biological replicates from the control, BS-N, PBS, and gas-phase conditions, and the bars indicate the mean values. Asterisks indicate statistical significance based on adjusted *p* values obtained by Dunnett’s test (* *p* < 0.05, ** *p* < 0.01, *** *p* < 0.001, **** *p* < 0.0001), with the control condition as the reference. For gene expression, each box represents a gene, and the color gradient indicates the log_2_ FC of gene expression in the BS-N, PBS, and gas-phase conditions relative to the control condition. Asterisks within the boxes indicate statistical significance based on the FDR calculated in the differential gene expression analysis (* FDR < 0.05, ** FDR < 0.01, *** FDR < 0.001). Abbreviations: TG, triacylglycerol; DG, diacylglycerol; MG, monoacylglycerol; FA, fatty acid; PA, phosphatidic acid; CDP-DG, cytidine diphosphate diacylglycerol; PS, phosphatidylserine; PGP, phosphatidylglycerophosphate; PE, phosphatidylethanolamine; PG, phosphatidylglycerol; LPE, lysophosphatidylethanolamine; CL, cardiolipin; MLCL, monolysocardiolipin; HBMP, hemibis(monoacylglycero)phosphate; CoQ, coenzyme Q; Glycerol-3P, glycerol 3-phosphate.

The two major storage lipids in Tol 5, WE and TG (Inoue et al., 2026), showed different patterns across the conditions. WE was detected at high levels only in the control cells and decreased in all three toluene degradation conditions. Consistently, the aldehyde dehydrogenase gene *calB*, predicted to participate in fatty alcohol assimilation (Inoue et al., 2026), was highly upregulated in the PBS and gas-phase conditions (Fig. 6). In contrast, TG remained abundant in BS-N and PBS but decreased only in the gas phase (Fig. 6). The levels of DG and MG, produced by TG degradation, also decreased in this condition. Among the genes annotated as lipases in the Tol 5 genome, TOL5_26490, TOL5_29350, and TOL5_29360 were specifically upregulated in the gas phase (Table S5). Degradation of these storage lipids releases fatty acids, and the genes *fadD* and *fadD4*, encoding fatty acyl-CoA ligases that activate fatty acids for β-oxidation, were more highly expressed in the gas phase than in the other conditions (Fig. 6). MG degradation also releases glycerol, and the genes *glpK* and *glpD*, which mediate glycerol utilization, were upregulated in the gas phase (Fig. 6). Glycerol utilization yields dihydroxyacetone phosphate, which is converted to glyceraldehyde 3-phosphate and enters glycolysis and gluconeogenesis.

Both membrane phospholipids PE and PG increased along with their precursor PA in the gas phase (Fig. 6). The species composed of 16:0 and 18:1 increased most markedly in PE and PG, whereas the increase in PA was dominated by saturated species (Fig. S5). Because the saturated PG species (16:0_16:0 and 16:0_18:0) were unchanged, the increase in the 16:0_18:1 species raised the proportion of unsaturated acyl chains in PG. Among the desaturase genes in Tol 5, TOL5_25690 was highly upregulated in the gas phase (Table S5). Despite the increased PE and PG, genes converting PA into PE and PG showed a decreasing trend. Among the membrane lipids derived from PG and PE, the tri- and tetraacylated species (CL, MLCL, and HBMP) did not change across conditions, whereas LPE increased in all three toluene degradation conditions.

The CoQ species, which mediate electron transfer in the respiratory chain, contributed strongly to the separation of the gas-phase samples along PC1 (Fig. 3C), and all three detected species, CoQ7, CoQ8, and CoQ9, were elevated in the gas phase (Fig. 6). Within the ubiquinol biosynthesis pathway, only 3-demethoxyubiquinol 3-hydroxylase (*coq7*) was upregulated in the gas phase, while the expression of the other genes did not change across conditions (Table S5).

## 4 Discussion

Bacterial metabolism has generally been studied under growth conditions in liquid cultures or on agar media. However, bacteria in natural environments reside in more complex and heterogeneous habitats, where they are often subjected to physicochemical conditions distinct from those in aqueous media (Chan et al., 2012; Tecon and Or, 2017; Bryan et al., 2019; Flemming and Wuertz, 2019; Grinberg et al., 2019). Recent research in environmental microbiology and bioprocess engineering has increasingly focused on the cellular functions of microorganisms in gaseous environments (Shen et al., 2017; Grinberg et al., 2019; Usami et al., 2020; Oswin Henry et al., 2023; Xu et al., 2023). In this study, we conducted an integrated analysis of the metabolome, lipidome, and transcriptome of *Acinetobacter* sp. Tol 5 degrading toluene on a solid surface in the absence of bulk water, to elucidate its metabolic alterations. The results revealed that intracellular metabolism was reorganized, specifically involving alterations in nitrogen and lipid metabolism (Fig. 7). These responses likely represent an adaptation strategy that enables the cells to maintain metabolic activity even in the gas phase. Previous studies on bacterial physiology in gaseous environments have primarily focused on desiccation tolerance, typically reporting a decrease in metabolic activity and a transition to a dormant state (Lebre et al., 2017; Pazos-Rojas et al., 2019; Lucidi et al., 2025). This study is distinct in that it analyzed metabolically active cells utilizing a gaseous carbon source, toluene, offering new insights into the survival strategies of microorganisms that utilize volatile substrates in natural environments.

**Figure 7.**
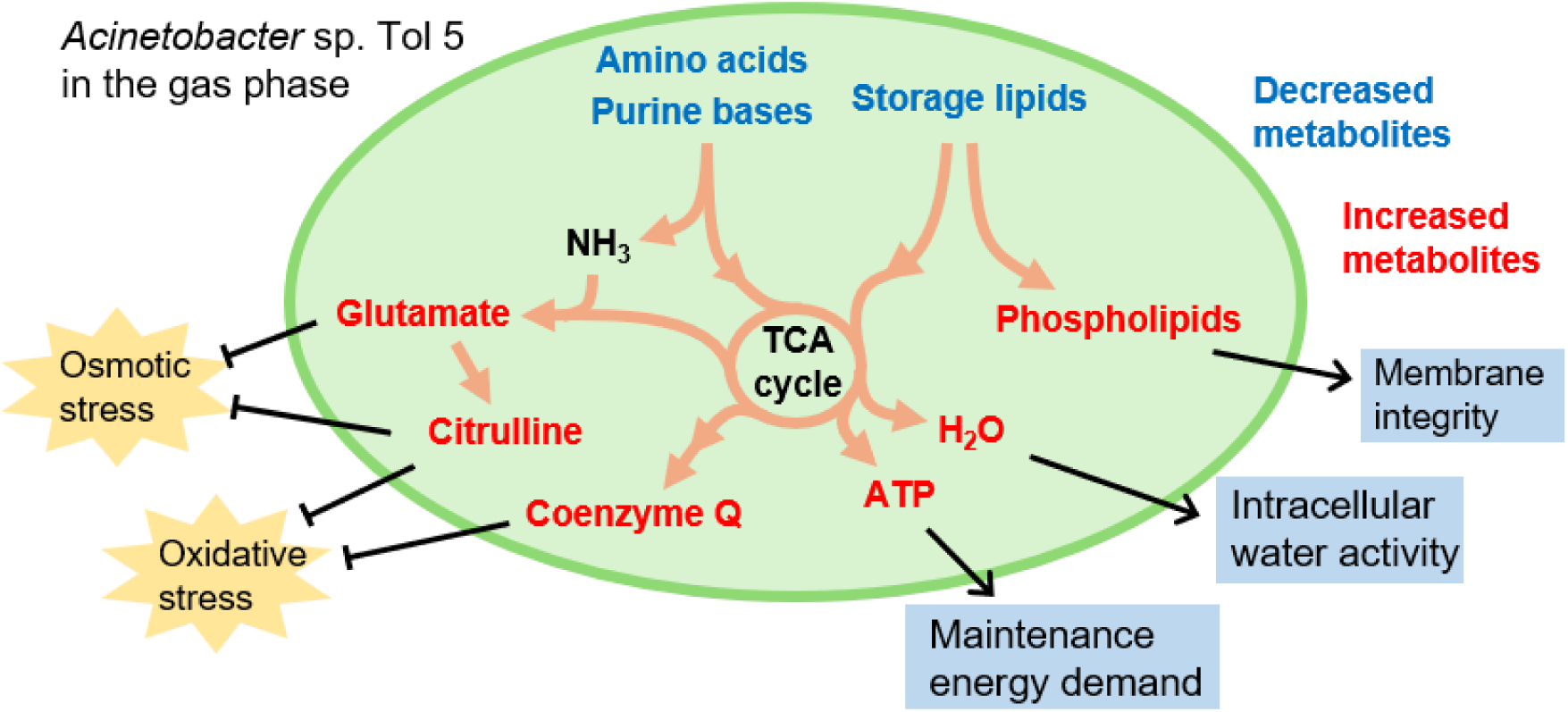
Schematic overview of metabolic alterations in *Acinetobacter* sp. Tol 5 under the gas-phase condition.

The most prominent difference between the aqueous and gas-phase conditions was in nitrogen metabolism. Amino acid and nucleobase degradation was enhanced in the gas phase, whereas glutamate and citrulline accumulated. Because external nitrogen was absent in all three conditions, these results indicate a distinct nitrogen starvation response specific to the gas-phase condition. Glutamate is the principal amino-group donor in bacterial nitrogen metabolism (Yan, 2007; Schulz-Mirbach et al., 2022) and also serves as a compatible solute and counterion for intracellular potassium (Dinnbier et al., 1988; Sleator and Hill, 2002). In *Acinetobacter baumannii*, glutamate accumulates under desiccation (Zeidler and Müller, 2019). Its maintenance in Tol 5 despite nitrogen limitation may therefore reflect a dual demand, for nitrogen donation and for osmotic adjustment in the gas phase. Citrulline is less well characterized in bacteria. It accumulates under salt stress in osmotolerant suppressor mutants derived from proline-biosynthesis-defective *Bacillus subtilis* strains, where it has been proposed to act as an alternative compatible solute (Stecker et al., 2022). Furthermore, in plants, particularly members of the Cucurbitaceae, citrulline accumulates under severe drought stress and acts as a hydroxyl radical scavenger that protects DNA and proteins (Akashi et al., 2001; Song et al., 2020). Given that the gas phase combines limited water availability with potential oxidative stress from the high oxygen availability, citrulline accumulation in Tol 5 may be functionally analogous.

In addition to these metabolite changes, genes encoding the enzymes of GS–GOGAT cycle, a major ammonia assimilation pathway in Gram-negative bacteria, were upregulated in the gas phase. The *nas* genes for assimilatory nitrate reduction were also upregulated, even though no nitrate was present in any condition. Because these genes are typically controlled by global nitrogen regulatory systems, their induction indicates that the cells sensed nitrogen limitation most strongly in the gas phase. Consistent with this, genes for aromatic amino acid catabolism and the degradation pathways for both purine and pyrimidine bases were induced most strongly in the gas phase. In *Escherichia coli*, genes involved in pyrimidine degradation are induced at the onset of nitrogen starvation, whereas degradation of allantoin, a purine catabolite, contributes to adaptation during prolonged starvation (Switzer et al., 2020). The concurrent activation of the purine and pyrimidine degradation pathways in Tol 5 suggests a stronger starvation response in the gas phase.

This study newly determined the lipid composition of Tol 5, which consists primarily of PE and PG, as in typical Gram-negative bacteria (Walczak-Skierska et al., 2024). Triacylated species, MLCL and acyl-PG (annotated as HBMP), were also abundant in Tol 5; both have been reported in other *Acinetobacter* strains (Lopalco et al., 2017; Luo et al., 2018). In *Acinetobacter radioresistens,* MLCL increases in relative abundance during stationary phase and has been linked to nutrient starvation (Luo et al., 2018), whereas in Tol 5 neither MLCL nor acyl-PG changed significantly across conditions, suggesting more intricate regulation of these triacylated species that warrants further study. The membrane phospholipids were remodeled in two respects with potentially opposing effects on fluidity. The proportions of PE and PG increased, and the increase in PE could contribute to reduced fluidity, as shown in liposome studies (Dawaliby et al., 2016). This interpretation is consistent with the reduced lateral diffusion inferred from the lipid ontology analysis in this study. Lower lateral diffusion generally reduces membrane permeability, which in a water-limited environment could limit the efflux of small molecules (Lande et al., 1995). Conversely, the selective enrichment of 18:1-containing species observed in the gas-phase condition would tend to increase fluidity by preventing a transition to a rigid gel phase (Yoon et al., 2015). These changes suggest that membrane properties were adjusted through two counteracting modifications rather than shifted unidirectionally. CoQ, which functions both as a respiratory electron carrier and as a membrane antioxidant (Søballe and Poole, 2000; Aussel et al., 2014), also increased in the gas phase. In *E. coli*, ubiquinone has been implicated in protection against the oxidative stress associated with long-chain fatty acid degradation (Agrawal et al., 2017); given the concurrent enhancement of β-oxidation in Tol 5, the increased CoQ may contribute to mitigating oxidative stress from this catabolic activity and from gas-phase exposure.

Many environmental bacteria accumulate intracellular WE and TG as energy storage compounds when carbon sources are abundant (Alvarez, 2016; Martin et al., 2021). Notably, *Acinetobacter* species exhibit a high accumulation capacity, with multiple wild-type strains accumulating WEs to over 10% of their cellular dry weight (Martin et al., 2021). In this study, WE decreased in all three toluene degradation conditions, whereas TG decreased only in the gas phase. The upregulation of genes encoding fatty acyl-CoA ligase (*fadD4*) and glycerol kinase (*glpK*) is consistent with enhanced catabolism of these storage lipids in the gas-phase condition (Fig. 6). Fatty acids released from storage lipids enter central metabolism via β-oxidation, and the glycerol moiety of TG enters glycolysis and gluconeogenesis. Consistent with this, glycolytic and gluconeogenic intermediates were among the metabolites distinguishing the gas-phase cells (Fig. 2C), although the present data cannot distinguish carbon derived from storage lipids from that derived from toluene. Storage lipid consumption may thus contribute to meeting an increased energy demand in the gas phase or to supplying precursors for membrane remodeling. Beyond providing energy, the oxidative degradation of WE and TG yields metabolic water, with a higher yield per unit mass than that from carbohydrates (Frank, 1988; Röttig et al., 2016). In some desert-dwelling bacteria, metabolic water from storage lipid degradation has been proposed to contribute to desiccation tolerance (Röttig et al., 2016). The gas-phase-specific consumption of TG is therefore consistent with a role for storage lipids as an endogenous source of both energy and water when extracellular water is limited. This suggests that lipid storage capacity is a relevant trait for strain selection. In this regard, *Acinetobacter* strains with a high capacity for storage lipid accumulation (Alvarez, 2016) are promising chassis for gas-phase bioprocesses.

In the gas-phase bioprocess examined here, water was partially retained in the porous carriers, and the immobilized cells functioned in low-water environments without a surrounding bulk aqueous medium. Their metabolic state differed markedly from that of cells growing on toluene in nitrogen-replete BS medium and was also distinct from that of resting cells degrading toluene in nitrogen-free aqueous media (BS-N and PBS). These findings identify metabolic features associated with gas-phase operation beyond those accompanying the transition from growth to resting-cell conditions, providing a basis for understanding cellular adaptation and for designing robust gas-phase bioprocesses.

## 5 Conclusion

This study characterized the metabolic alterations in metabolically active Tol 5 cells degrading toluene in the aqueous and gas phases. Integrated omics analysis showed that the gas-phase cells maintained their glutamate pool and exhibited enhanced amino acid and nucleobase degradation, suggesting endogenous nitrogen mobilization. The gas-phase-specific consumption of the storage lipid triacylglycerol suggests a role for this lipid as an endogenous source of energy and, potentially, metabolic water. These findings identify nitrogen availability and lipid storage capacity as considerations for the rational design of gas-phase bioprocesses and for the selection of microbial chassis.

## CRediT authorship contribution statement

**Shori Inoue:** Conceptualization, Formal analysis, Investigation, Data curation, Writing - Original Draft. **Taisei Naobayashi:** Investigation, Data curation, Writing - Original Draft. **Kanako Tokiyoshi:** Investigation, Data curation, Writing - Review & Editing. **Shogo Yoshimoto:** Writing - Review & Editing. **Hiroshi Tsugawa:** Conceptualization, Supervision, Writing - Review & Editing. **Katsutoshi Hori:** Conceptualization, Supervision, Project administration, Writing - Review & Editing.

## Data availability

All raw MS data are available in the MB-POST repository (https://repository.massbank.jp/) under index number MPST000146. Although the processed table data used to create the figures were submitted as source data, the original peak lists and alignment tables were deposited in MB-POST together with the raw data.

RNA-seq data reported are available in the DDBJ Sequenced Read Archive under the accession number PRJDB43029.

## Acknowledgements

The authors wish to acknowledge the Center for Gene Research, Nagoya University, for technical support with the RNA sequencing.

This research was supported by the Graduate Program of Transformative Chem-Bio Research at Nagoya University supported by MEXT (WISE Program) to SI, the Japan Science and Technology Agency (JST) SPRING (Grant Number JPMJSP2125) to SI, the GteX Program Japan (Grant Number JPMJGX23B4) to KH, the JST Exploratory Research for Advanced Technology (ERATO) (Grant Number JPMJER2101) to HT, the JST FOREST Program (Grant Number JPMJFR230H) to HT, the JST NBDC Program (Grant Number JPMJND2305) to HT, and the Japan Society for the Promotion of Science (JSPS) KAKENHI (Grant Numbers JP24H00043 and JP25K24646 to KH, HT, and SY; JP26K01314 to SY and KH; and JP24K02011, JP24H00392, JP24K21269, JP25H01425, and JP25H01426 to HT).

## References

1. Agrawal S, Jaswal K, Shiver AL, Balecha H, Patra T, Chaba R. A genome-wide screen in *Escherichia coli* reveals that ubiquinone is a key antioxidant for metabolism of long-chain fatty acids. Journal of Biological Chemistry 2017;292(49):20086–20099.

2. Akashi K, Miyake C, Yokota A. Citrulline, a novel compatible solute in drought-tolerant wild watermelon leaves, is an efficient hydroxyl radical scavenger. FEBS Letters 2001;508(3):438–442.

3. Alvarez HM. Triacylglycerol and wax ester-accumulating machinery in prokaryotes. Biochimie 2016;120:28–39.

4. Asimakopoulos K, Gavala HN, Skiadas IV. Reactor systems for syngas fermentation processes: A review. Chemical Engineering Journal 2018;348:732–744.

5. Aussel L, Pierrel F, Loiseau L, Lombard M, Fontecave M, Barras F. Biosynthesis and physiology of coenzyme Q in bacteria. Biochimica et Biophysica Acta (BBA) - Bioenergetics 2014;1837(7):1004–1011.

6. Bligh EG, Dyer WJ. A rapid method of total lipid extraction and purification. Canadian Journal of Biochemistry and Physiology 1959;37(8):911–7.

7. Brown GR, Sutcliffe IC, Cummings SP. Combined solvent and water activity stresses on turgor regulation and membrane adaptation in *Oceanimonas baumannii* ATCC 700832. Antonie van Leeuwenhoek 2003;83(3):275–283.

8. Bryan NC, Christner BC, Guzik TG, Granger DJ, Stewart MF. Abundance and survival of microbial aerosols in the troposphere and stratosphere. The ISME Journal 2019;13(11):2789–2799.

9. Chan Y, Lacap DC, Lau MCY, Ha KY, Warren-Rhodes KA, Cockell CS, Cowan DA, McKay CP, Pointing SB. Hypolithic microbial communities: between a rock and a hard place. Environmental Microbiology 2012;14(9):2272–2282.

10. Chen S, Zhou Y, Chen Y, Gu J. fastp: an ultra-fast all-in-one FASTQ preprocessor. Bioinformatics 2018;34(17):i884–i890.

11. Chen Y-Y, Ishikawa M, Hori K. A novel inverse membrane bioreactor for efficient bioconversion from methane gas to liquid methanol using a microbial gas-phase reaction. Biotechnology for Biofuels and Bioproducts 2023;16(1):16.

12. Dawaliby R, Trubbia C, Delporte C, Noyon C, Ruysschaert J-M, Van Antwerpen P, Govaerts C. Phosphatidylethanolamine Is a Key Regulator of Membrane Fluidity in Eukaryotic Cells*. Journal of Biological Chemistry 2016;291(7):3658–3667.

13. Devi NB, Pakshirajan K. Diversifying product portfolio of syngas fermentation in addition to ethanol production by using *Clostridium species*. Bioresource Technology 2025;427:132401.

14. Dinnbier U, Limpinsel E, Schmid R, Bakker EP. Transient accumulation of potassium glutamate and its replacement by trehalose during adaptation of growing cells of *Escherichia coli* K-12 to elevated sodium chloride concentrations. Archives of Microbiology 1988;150(4):348–357.

15. Flemming HC, Wuertz S. Bacteria and archaea on Earth and their abundance in biofilms. Nature Reviews Microbiology 2019;17(4):247–260.

16. Frank CL. Diet Selection by a Heteromyid Rodent: Role of Net Metabolic Water Production. Ecology 1988;69(6):1943–1951.

17. Grinberg M, Orevi T, Steinberg S, Kashtan N. Bacterial survival in microscopic surface wetness. eLife 2019;8:e48508.

18. Hori K, Yamashita S, Ishii S, Kitagawa M, Tanji Y, Unno H. Isolation, Characterization and Application to Off-Gas Treatment of Toluene-Degrading Bacteria. Journal of Chemical Engineering of Japan 2001;34(9):1120–1126.

19. Imlay JA. The molecular mechanisms and physiological consequences of oxidative stress: lessons from a model bacterium. Nature Reviews Microbiology 2013;11(7):443–454.

20. Inoue S, Yoshimoto S, Hattori M, Yamagishi S, Hori K. Characterization of carbon metabolism in a highly adhesive bacterium *Acinetobacter* sp. Tol 5 capable of assimilating diverse hydrocarbons and aromatic compounds. Frontiers in Microbiology 2026;17:1919423.

21. Ishikawa M, Nakatani H, Hori K. AtaA, a New Member of the Trimeric Autotransporter Adhesins from *Acinetobacter* sp. Tol 5 Mediating High Adhesiveness to Various Abiotic Surfaces. PLOS ONE 2012;7(11):e48830.

22. Jawad A, Seifert H, Snelling AM, Heritage J, Hawkey PM. Survival of *Acinetobacter baumannii* on Dry Surfaces: Comparison of Outbreak and Sporadic Isolates. Journal of Clinical Microbiology 1998;36(7):1938–1941.

23. Kiuchi S, Otoguro Y, Nitta T, Chung MH, Nakaya T, Matsuzawa Y, Ohbuchi K, Sasaki K, Yamamoto H, Tsugawa H. Using Variable Data-Independent Acquisition for Capillary Electrophoresis-Based Untargeted Metabolomics. Journal of the American Society for Mass Spectrometry 2024;35(9):2118–2127.

24. Klåvus A, Kokla M, Noerman S, Koistinen VM, Tuomainen M, Zarei I, Meuronen T, Häkkinen MR, Rummukainen S, Farizah Babu A, Sallinen T, Kärkkäinen O, Paananen J, Broadhurst D, Brunius C, Hanhineva K. “notame”: Workflow for Non-Targeted LC-MS Metabolic Profiling. Metabolites 2020;10(4):135.

25. Lande MB, Donovan JM, Zeidel ML. The relationship between membrane fluidity and permeabilities to water, solutes, ammonia, and protons. Journal of General Physiology 1995;106(1):67–84.

26. Langmead B, Salzberg SL. Fast gapped-read alignment with Bowtie 2. Nature Methods 2012;9(4):357–359.

27. Lebre PH, De Maayer P, Cowan DA. Xerotolerant bacteria: surviving through a dry spell. Nature Reviews Microbiology 2017;15(5):285–296.

28. Liao Y, Smyth GK, Shi W. featureCounts: an efficient general purpose program for assigning sequence reads to genomic features. Bioinformatics 2014;30(7):923–930.

29. Liew FE, Nogle R, Abdalla T, Rasor BJ, Canter C, Jensen RO, Wang L, Strutz J, Chirania P, De Tissera S, Mueller AP, Ruan Z, Gao A, Tran L, Engle NL, Bromley JC, Daniell J, Conrado R, Tschaplinski TJ, Giannone RJ, Hettich RL, Karim AS, Simpson SD, Brown SD, Leang C, Jewett MC, Köpke M. Carbon-negative production of acetone and isopropanol by gas fermentation at industrial pilot scale. Nature Biotechnology 2022;40(3):335–344.

30. Lopalco P, Stahl J, Annese C, Averhoff B, Corcelli A. Identification of unique cardiolipin and monolysocardiolipin species in *Acinetobacter baumannii*. Scientific Reports 2017;7(1):2972.

31. Lopes M, Brejchova K, Riecan M, Novakova M, Rossmeisl M, Cajka T, Kuda O. Metabolomics atlas of oral _13_C-glucose tolerance test in mice. Cell Reports 2021;37(2):109833.

32. Lucidi M, Capecchi G, Spagnoli C, Basile A, Artuso I, Persichetti L, Fardelli E, Capellini G, Visaggio D, Imperi F, Rampioni G, Leoni L, Visca P. The response to desiccation in *Acinetobacter baumannii*. Virulence 2025;16(1):2490209.

33. Luo Y, Javed MA, Deneer H, Chen X. Nutrient depletion-induced production of tri-acylated glycerophospholipids in *Acinetobacter radioresistens*. Scientific Reports 2018;8(1):7470.

34. Martin LK, Huang WE, Thompson IP. Bacterial wax synthesis. Biotechnology Advances 2021;46:107680.

35. Molenaar MR, Jeucken A, Wassenaar TA, van de Lest CHA, Brouwers JF, Helms JB. LION/web: a web-based ontology enrichment tool for lipidomic data analysis. Gigascience 2019;8(6):giz061.

36. Morono Y, Unno H, Tanji Y, Hori K. Addition of Aromatic Substrates Restores Trichloroethylene Degradation Activity in *Pseudomonas putida* F1. Applied and Environmental Microbiology 2004;70(5):2830–2835.

37. Neto AS, Wainaina S, Chandolias K, Piatek P, Taherzadeh MJ. Exploring the Potential of Syngas Fermentation for Recovery of High-Value Resources: A Comprehensive Review. Current Pollution Reports 2024;11(1):7.

38. Ohara Y, Yoshimoto S, Hori K. Control of AtaA-mediated bacterial immobilization by casein hydrolysates. Journal of Bioscience and Bioengineering 2019;128(5):544–550.

39. Oswin Henry P, Haddrell Allen E, Hughes C, Otero-Fernandez M, Thomas Richard J, Reid Jonathan P. Oxidative Stress Contributes to Bacterial Airborne Loss of Viability. Microbiology Spectrum 2023;11(2):e03347–22.

40. Pang Z, Lu Y, Zhou G, Hui F, Xu L, Viau C, Spigelman AF, MacDonald PE, Wishart DS, Li S, Xia J. MetaboAnalyst 6.0: towards a unified platform for metabolomics data processing, analysis and interpretation. Nucleic Acids Research 2024;52(W1):W398–W406.

41. Pazos-Rojas LA, Muñoz-Arenas LC, Rodríguez-Andrade O, López-Cruz LE, López-Ortega O, Lopes-Olivares F, Luna-Suarez S, Baez A, Morales-García YE, Quintero-Hernández V, Villalobos-López MA, De la Torre J, Muñoz-Rojas J. Desiccation-induced viable but nonculturable state in *Pseudomonas putida* KT2440, a survival strategy. PLOS ONE 2019;14(7):e0219554.

42. Romantsov T, Guan Z, Wood JM. Cardiolipin and the osmotic stress responses of bacteria. Biochimica et Biophysica Acta (BBA) - Biomembranes 2009;1788(10):2092–2100.

43. Röttig A, Hauschild P, Madkour MH, Al-Ansari AM, Almakishah NH, Steinbüchel A. Analysis and optimization of triacylglycerol synthesis in novel oleaginous *Rhodococcus* and *Streptomyces* strains isolated from desert soil. Journal of Biotechnology 2016;225:48–56.

44. Rybarczyk P, Szulczyński B, Gębicki J, Hupka J. Treatment of malodorous air in biotrickling filters: A review. Biochemical Engineering Journal 2019;141:146–162.

45. Schulz-Mirbach H, Müller A, Wu T, Pfister P, Aslan S, Schada von Borzyskowski L, Erb TJ, Bar-Even A, Lindner SN. On the flexibility of the cellular amination network in *E coli*. eLife 2022;11:e77492.

46. Schymanski EL, Jeon J, Gulde R, Fenner K, Ruff M, Singer HP, Hollender J. Identifying small molecules via high resolution mass spectrometry: communicating confidence. Environmental Science & Technology 2014;48(4):2097–8.

47. Shen Y, Brown RC, Wen Z. Syngas fermentation by *Clostridium carboxidivorans* P7 in a horizontal rotating packed bed biofilm reactor with enhanced ethanol production. Applied Energy 2017;187:585–594.

48. Sheoran K, Siwal SS, Kapoor D, Singh N, Saini AK, Alsanie WF, Thakur VK. Air Pollutants Removal Using Biofiltration Technique: A Challenge at the Frontiers of Sustainable Environment. ACS Engineering Au 2022;2(5):378–396.

49. Sleator RDHill C. Bacterial osmoadaptation: the role of osmolytes in bacterial stress and virulence. FEMS Microbiology Reviews 2002;26(1):49–71.

50. Søballe BPoole RK. Ubiquinone limits oxidative stress in *Escherichia coli*. Microbiology 2000;146(4):787–796.

51. Song Q, Joshi M, DiPiazza J, Joshi V. Functional Relevance of Citrulline in the Vegetative Tissues of Watermelon During Abiotic Stresses. Frontiers in Plant Science 2020;11:512.

52. Stecker D, Hoffmann T, Link H, Commichau FM, Bremer E. L-Proline Synthesis Mutants of *Bacillus subtilis* Overcome Osmotic Sensitivity by Genetically Adapting L-Arginine Metabolism. Frontiers in Microbiology 2022;13:908304.

53. Switzer A, Burchell L, McQuail J, Wigneshweraraj S. The Adaptive Response to Long-Term Nitrogen Starvation in *Escherichia coli* Requires the Breakdown of Allantoin. Journal of Bacteriology 2020;202(17):10.1128/jb.00172-20.

54. Takeda H, Matsuzawa Y, Takeuchi M, Takahashi M, Nishida K, Harayama T, Todoroki Y, Shimizu K, Sakamoto N, Oka T, Maekawa M, Chung MH, Kurizaki Y, Kiuchi S, Tokiyoshi K, Buyantogtokh B, Kurata M, Kvasnička A, Takeda U, Uchino H, Hasegawa M, Miyamoto J, Tanabe K, Takeda S, Mori T, Kumakubo R, Tanaka T, Yoshino T, Okamoto M, Takahashi H, Arita M, Tsugawa H. MS-DIAL 5 multimodal mass spectrometry data mining unveils lipidome complexities. Nature Communications 2024;15(1):9903.

55. Takors R, Kopf M, Mampel J, Bluemke W, Blombach B, Eikmanns B, Bengelsdorf FR, Weuster-Botz D, Dürre P. Using gas mixtures of CO, CO_2_ and H_2_ as microbial substrates: the do’s and don’ts of successful technology transfer from laboratory to production scale. Microbial Biotechnology 2018;11(4):606–625.

56. Tan JN, Ratra K, Singer SW, Simmons BA, Goswami S, Awasthi D. Methane to bioproducts: unraveling the potential of methanotrophs for biomanufacturing. Current Opinion in Biotechnology 2024;90:103210.

57. Tecon R, Or D. Biophysical processes supporting the diversity of microbial life in soil. FEMS Microbiology Reviews 2017;41(5):599–623.

58. Tokiyoshi K, Matsuzawa Y, Takahashi M, Takeda H, Hasegawa M, Miyamoto J, Tsugawa H. Using Data-Dependent and -Independent Hybrid Acquisitions for Fast Liquid Chromatography-Based Untargeted Lipidomics. Analytical Chemistry 2024;96(3):991–996.

59. Tsugawa H, Ikeda K, Takahashi M, Satoh A, Mori Y, Uchino H, Okahashi N, Yamada Y, Tada I, Bonini P, Higashi Y, Okazaki Y, Zhou Z, Zhu ZJ, Koelmel J, Cajka T, Fiehn O, Saito K, Arita M. A lipidome atlas in MS-DIAL 4. Nature Biotechnology 2020;38(10):1159–1163.

60. Usami A, Ishikawa M, Hori K. Gas-phase bioproduction of a high-value-added monoterpenoid (*E*)-geranic acid by metabolically engineered *Acinetobacter* sp. Tol 5. Green Chemistry 2020;22(4):1258–1268.

61. Walczak-Skierska J, Ludwiczak A, Sibińska E, Pomastowski P. Environmental Influence on Bacterial Lipid Composition: Insights from Pathogenic and Probiotic Strains. ACS Omega 2024;9(36):37789–37801.

62. Watanabe H, Tanji Y, Unno H, Hori K. Rapid conversion of toluene by an *acinetobacter* sp. Tol 5 mutant showing monolayer adsorption to water-oil interface. Journal of Bioscience and Bioengineering 2008;106(3):226–230.

63. Xia Q, Yang J, Hu L, Zhao H, Lu Y. Biotransforming CO_2_ into valuable chemicals. Journal of Cleaner Production 2024;434:140185.

64. Xu C, Frigo-Vaz B, Goering J, Wang P. Gas-phase degradation of VOCs using supported bacteria biofilms. Biotechnology and Bioengineering 2023;120(5):1323–1333.

65. Yan D. Protection of the glutamate pool concentration in enteric bacteria. Proceedings of the National Academy of Sciences 2007;104(22):9475–9480.

66. Yoon Y, Lee H, Lee S, Kim S, Choi K-H. Membrane fluidity-related adaptive response mechanisms of foodborne bacterial pathogens under environmental stresses. Food Research International 2015;72:25–36.

67. Yoshimoto S, Ohara Y, Nakatani H, Hori K. Reversible bacterial immobilization based on the salt-dependent adhesion of the bacterionanofiber protein AtaA. Microbial Cell Factories 2017;16(1):123.

68. Yu G, Wang L-G, Han Y, He Q-Y. clusterProfiler: an R Package for Comparing Biological Themes Among Gene Clusters. OMICS: A Journal of Integrative Biology 2012;16(5):284–287.

69. Zeidler S, Müller V. The role of compatible solutes in desiccation resistance of *Acinetobacter baumannii*. MicrobiologyOpen 2019;8(5):e00740.

