## Supplementary Information for "Integrated omics analysis reveals reorganization of nitrogen and lipid metabolism in a toluene-degrading bacterium in the gas phase"

**Supplemental figures**

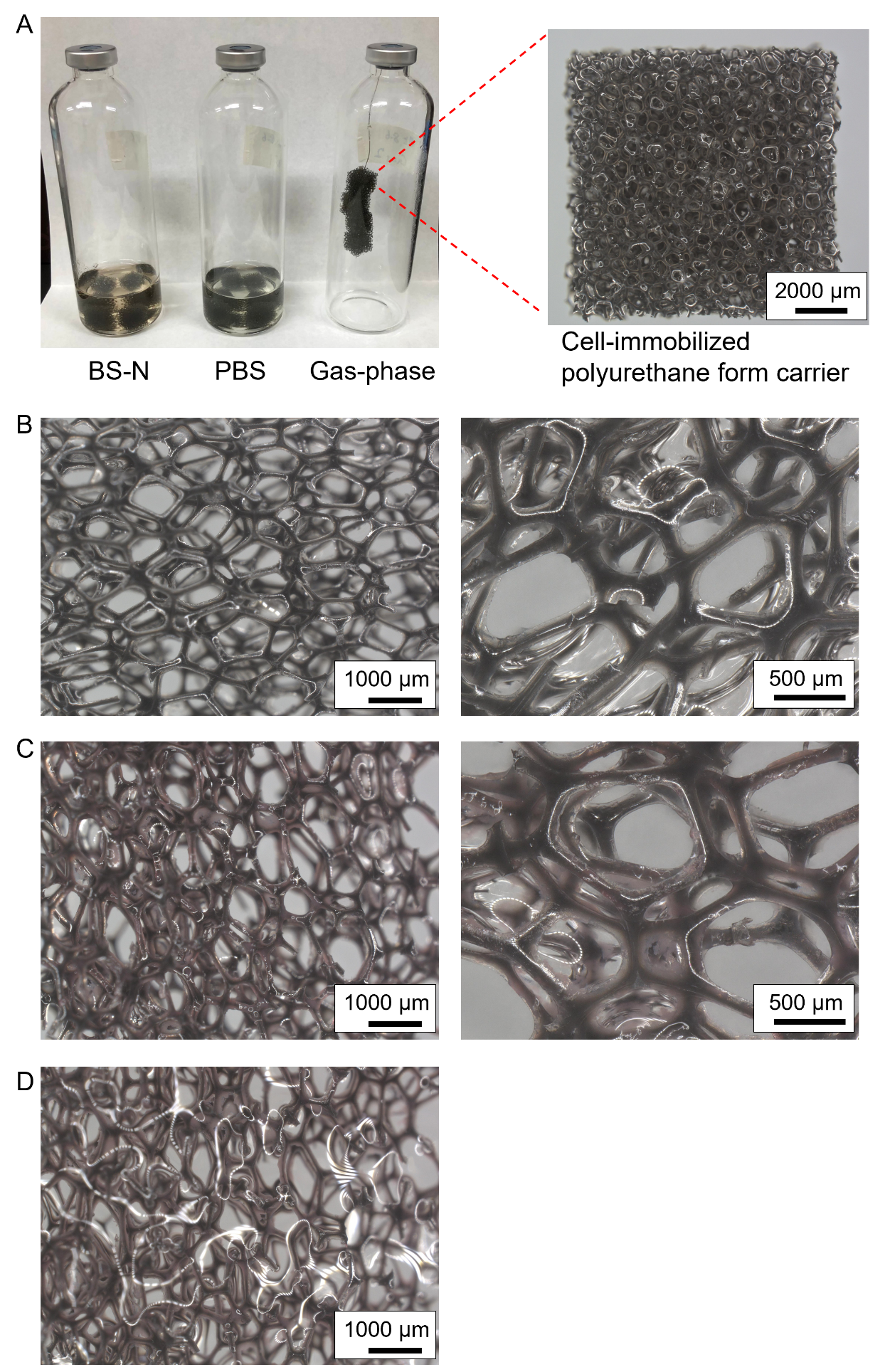

**Figure S1.** Experimental setup for the toluene degradation assay in the aqueous- and gas-phase conditions. (A) Vials for the BS-N, PBS, and gas-phase conditions. (B–D) Polyurethane foam without (B) or with (C, D) Tol 5 cells, visualized by propidium iodide staining after paraformaldehyde fixation. Images were taken after (B, C) or before (D) removal of excess liquid. Scale bars, 1000 µm (left) and 500 µm (right).

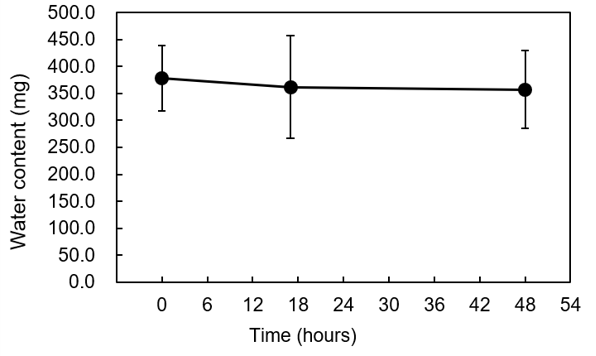

**Figure S2.** Water content of the PU carriers during the gas-phase reaction. The amount of water retained by each set of four carriers was determined gravimetrically from the difference in mass before and after lyophilization. Data are presented as mean ± standard deviation (n = 6 biological replicates).

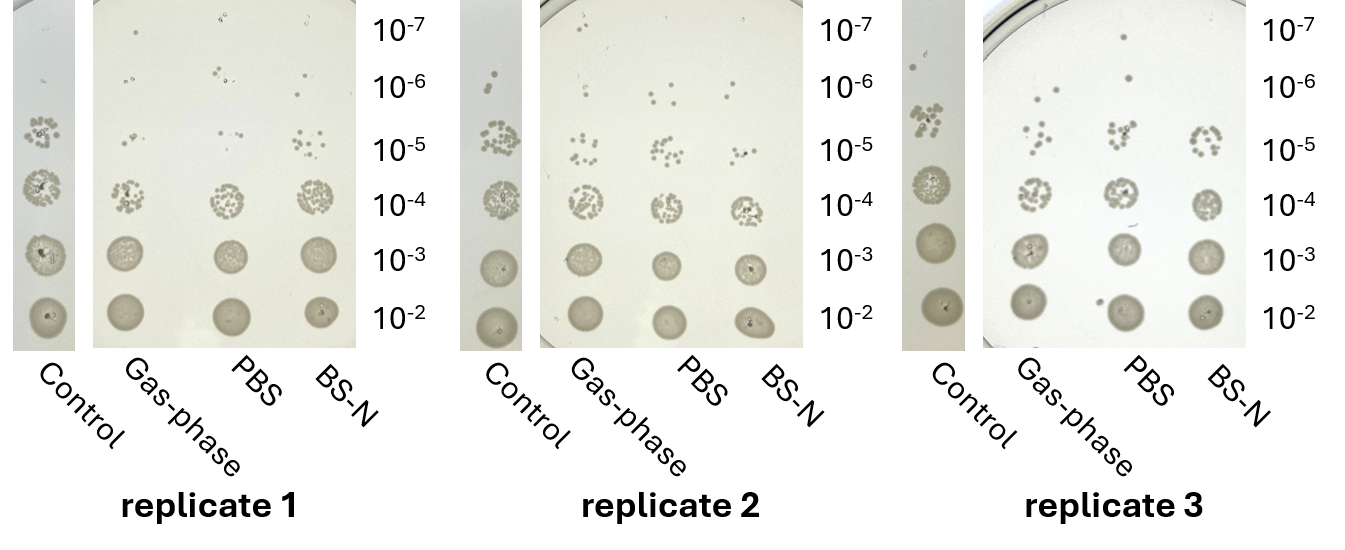

**Figure S3.** Colony formation of Tol 5 under the control, BS-N, PBS, and gas-phase conditions after a 48-hours toluene degradation assay. Cells before or after the toluene degradation assay were detached from the carriers, serially diluted, and inoculated onto LB agar plates.

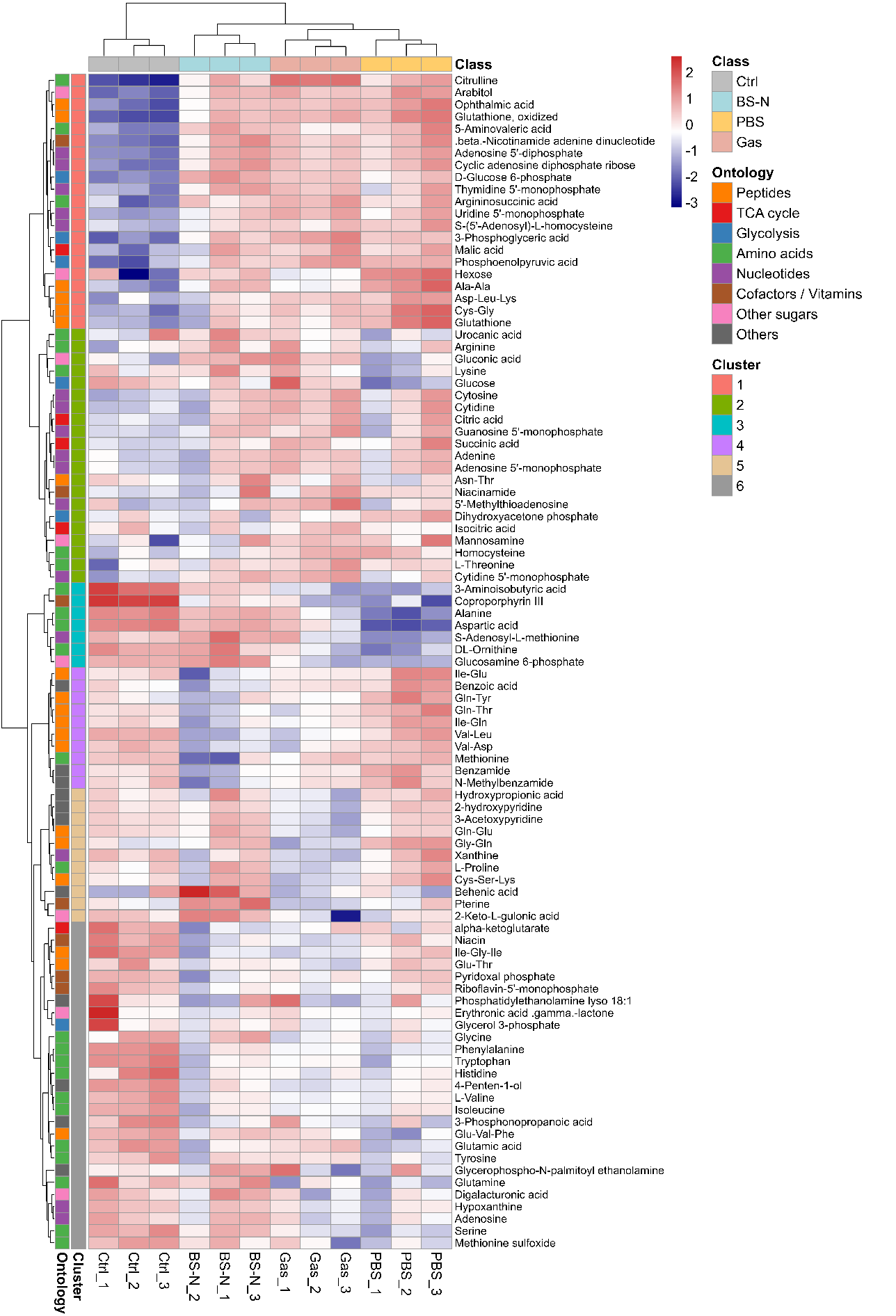

**Figure S4.** Hierarchical clustering analysis (HCA) of the metabolome data. The log₂-transformed values with a pseudo-count of 1 were scaled by the Pareto scaling method, with red and blue indicating high and low relative abundance, respectively.

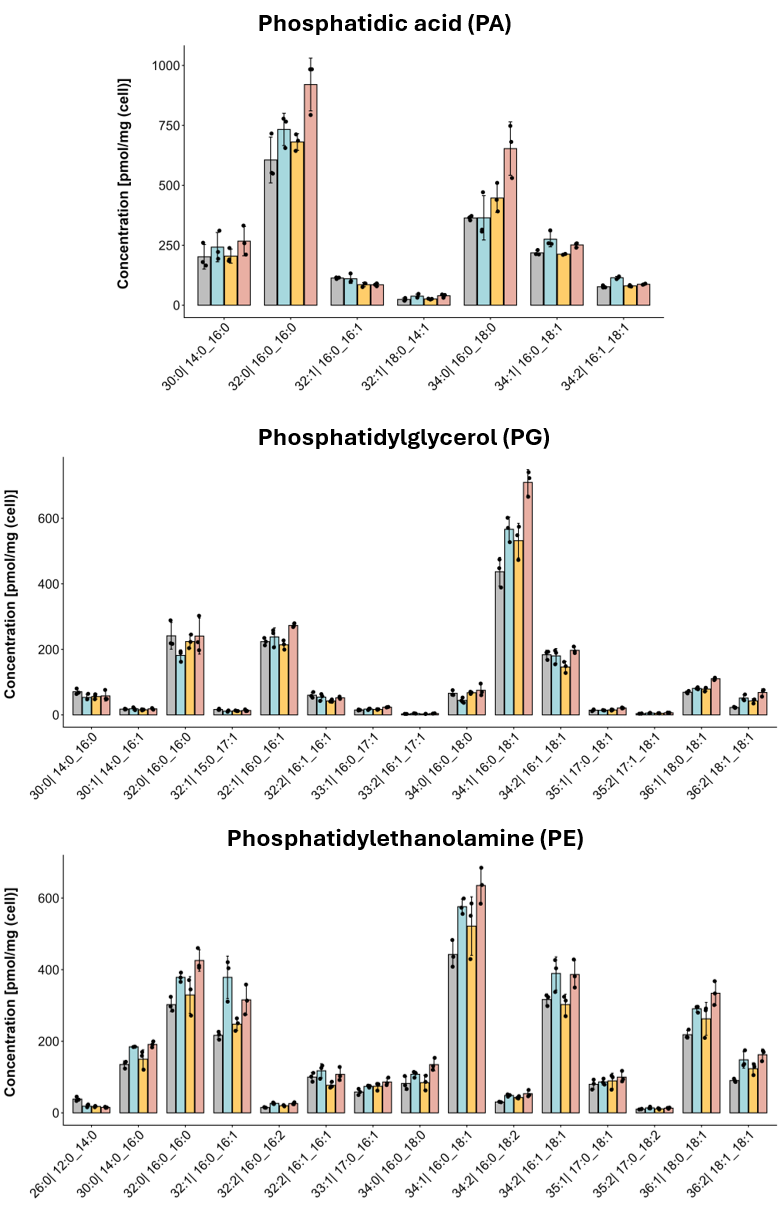

**Figure S5.** The composition of phosphatidic acid (PA), phosphatidylglycerol (PG), and phosphatidylethanolamine (PE). Each dot represents the concentration of three biological replicates from the control (gray), BS-N (light blue), PBS (orange), and gas-phase (pink) conditions, and the bars and error bars indicate the mean ± SD.

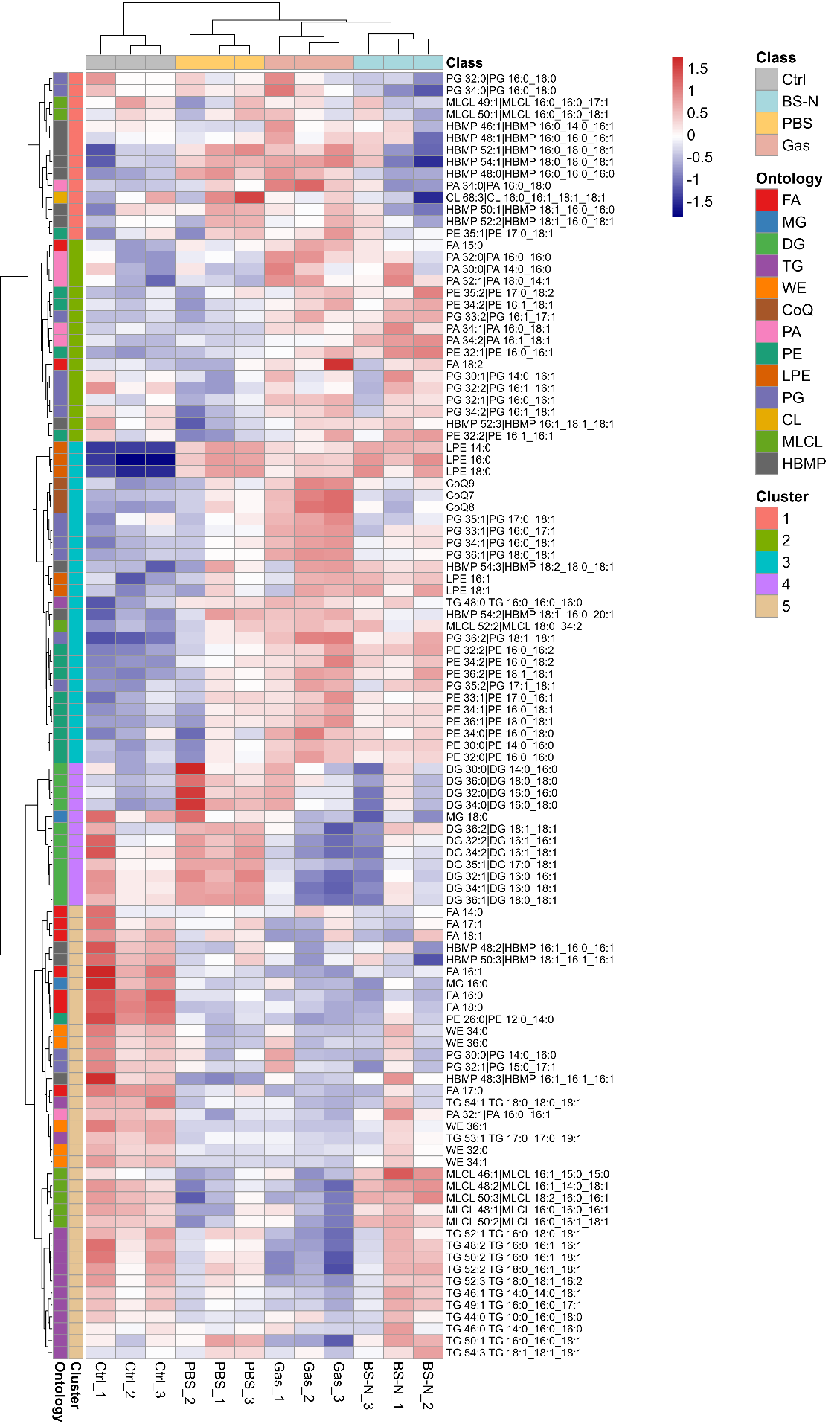

**Figure S6.** Hierarchical clustering analysis (HCA) of the lipidome data. The log₂-transformed values with a pseudo-count of 1 were scaled by the Pareto scaling method, with red and blue indicating high and low relative abundance, respectively.

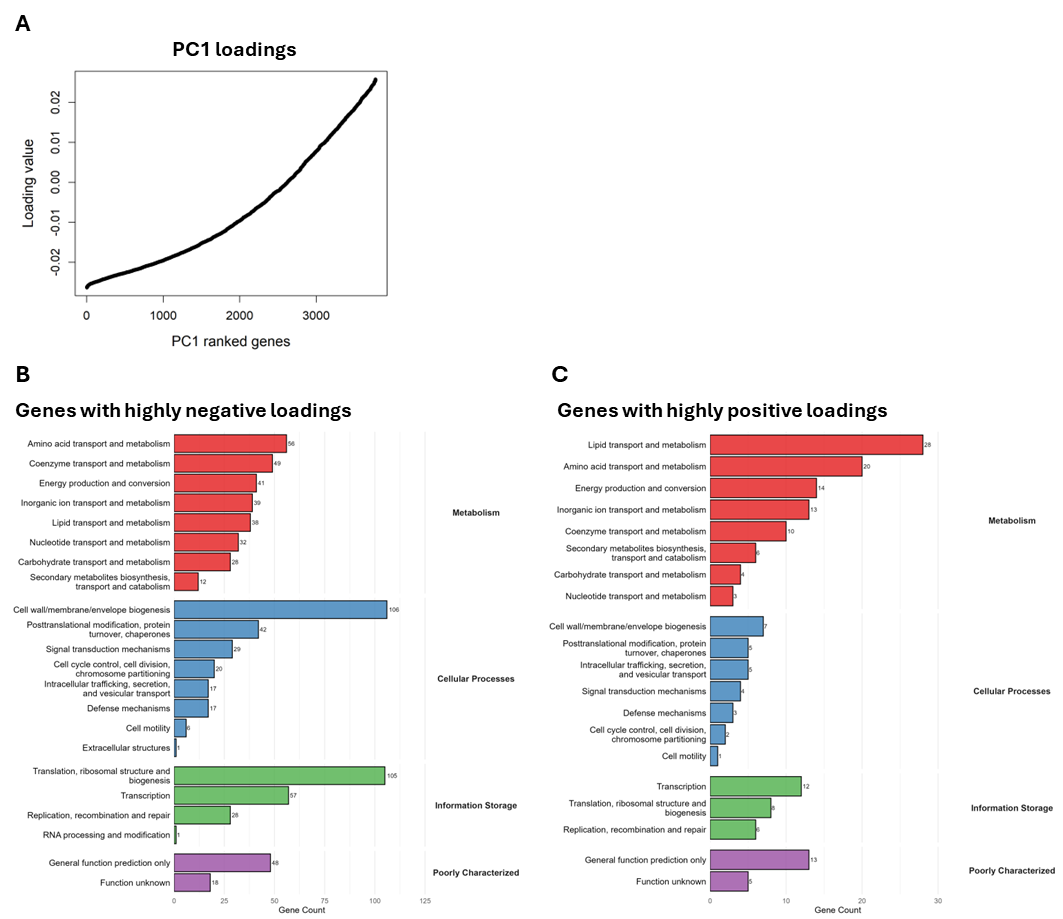

**Figure S7.** Distribution and functional classification of PC1 loading genes from the transcriptome principal component analysis shown in Fig. 4B. (**A**) Rank plot of PC1 loading values for all analyzed genes. Genes are ordered along the x-axis from the most negative to the most positive loading values. (**B**, **C**) Clusters of Orthologous Genes (COG) functional distribution of the genes with (**B**) highly negative (< -0.02) and (**C**) highly positive (> 0.02) PC1 loading values. The bar charts display the number of genes assigned to each COG category.

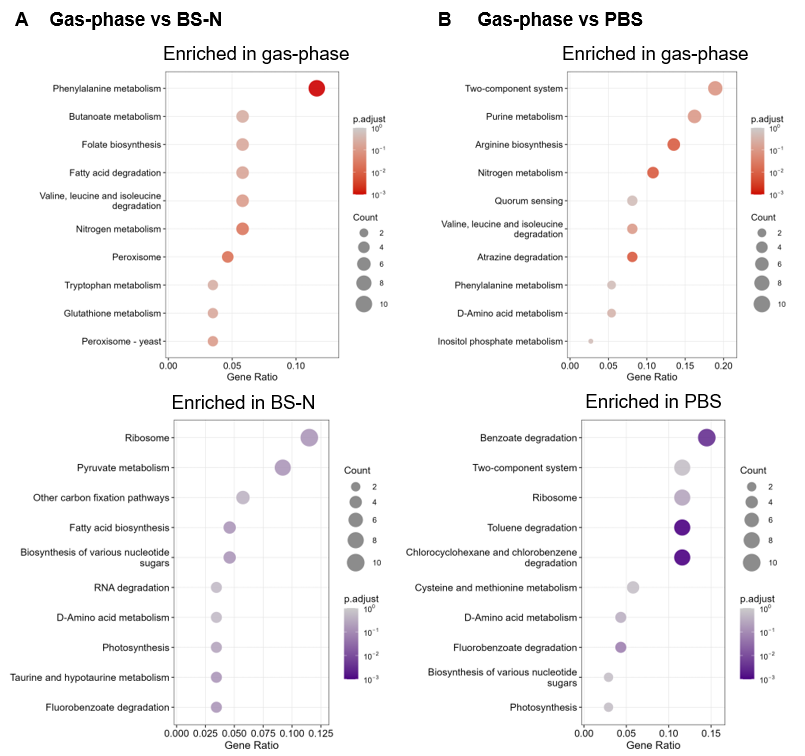

**Figure S8.** Over-representation analysis of differentially expressed genes across KEGG pathways, performed using clusterProfiler for (A) the gas phase compared with BS-N and (B) the gas phase compared with PBS. In each panel, pathways enriched among genes upregulated in the gas phase (upper) and among genes upregulated in the corresponding aqueous condition, i.e., downregulated in the gas phase (lower), are shown separately. The x-axis indicates the gene ratio, the dot size represents the number of differentially expressed genes assigned to each category, and the color gradient indicates the Benjamini–Hochberg adjusted *p* values.

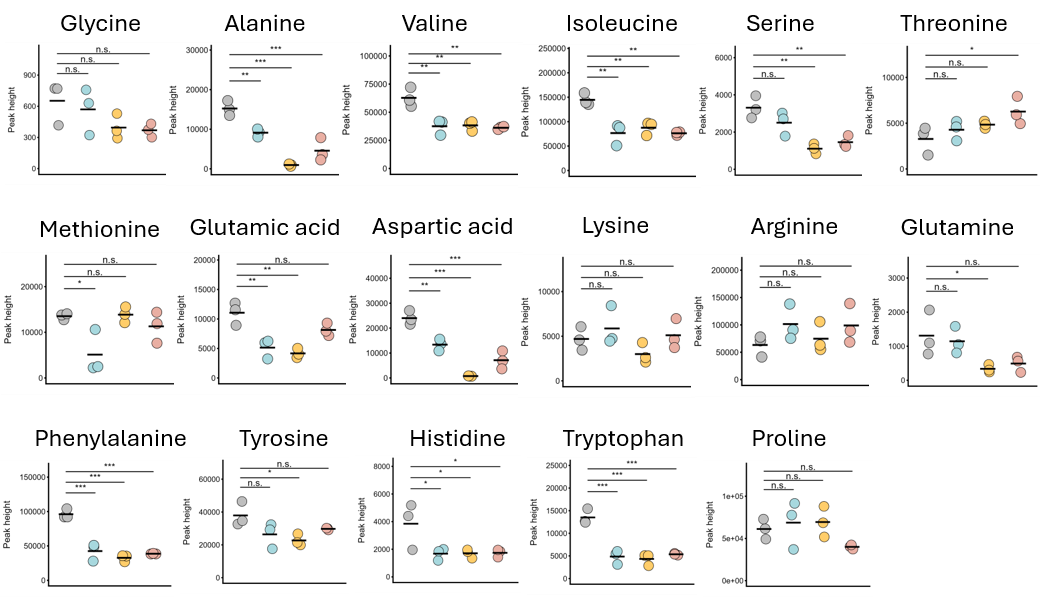

**Figure S9.** The levels of amino acids detected in the metabolome analysis. Each dot represents the normalized peak height of one of three biological replicates, and the bars indicate the mean values. Asterisks indicate statistical significance based on adjusted *p* values obtained by Dunnett's test (* *p* < 0.05, ** *p* < 0.01, *** *p* < 0.001, **** *p* < 0.0001), with the control condition as the reference.

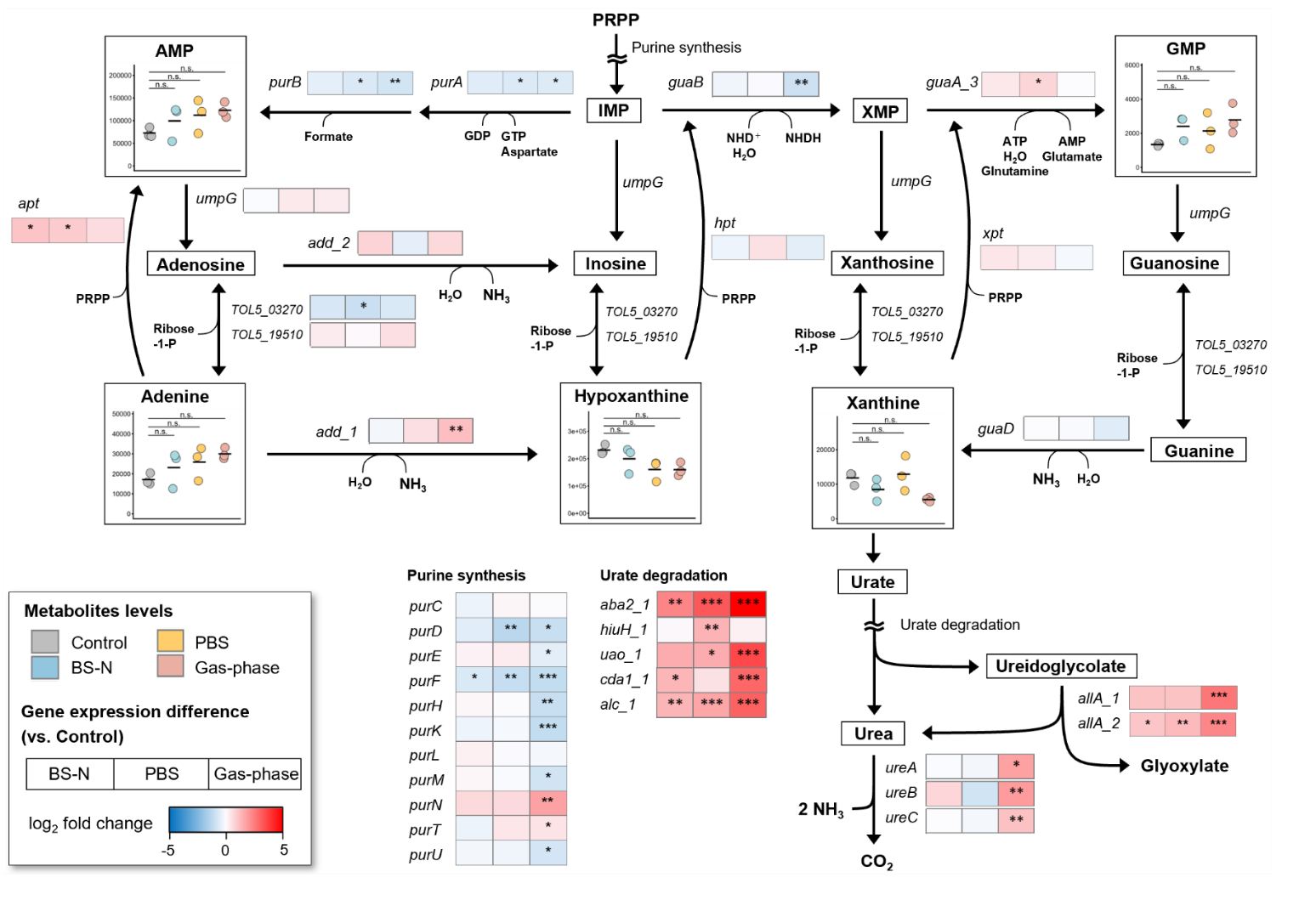

**Figure S10**. Metabolite levels and differential gene expression for purine metabolism. For metabolites, each dot represents the normalized peak height of one of three biological replicates from the control, BS-N, PBS, and gas-phase conditions, and the bars indicate the mean values. Asterisks indicate statistical significance based on adjusted *p* values obtained by Dunnett's test (* *p* < 0.05, ** *p* < 0.01, *** *p* < 0.001, **** *p* < 0.0001), with the control condition as the reference. For gene expression, each box represents a gene, and the color gradient indicates the log_2_ FC of gene expression in the BS-N, PBS, and gas-phase conditions relative to the control condition. Asterisks within the boxes indicate statistical significance based on the FDR calculated in the differential gene expression analysis (* FDR < 0.05, ** FDR < 0.01, *** FDR < 0.001). Abbreviations: PRPP, 5-phospho-α-D-ribose 1-diphosphate; IMP, inosine 5′-monophosphate; XMP, xanthosine 5′-monophosphate; AMP, adenosine 5′-monophosphate; GMP, guanosine 5′-monophosphate.

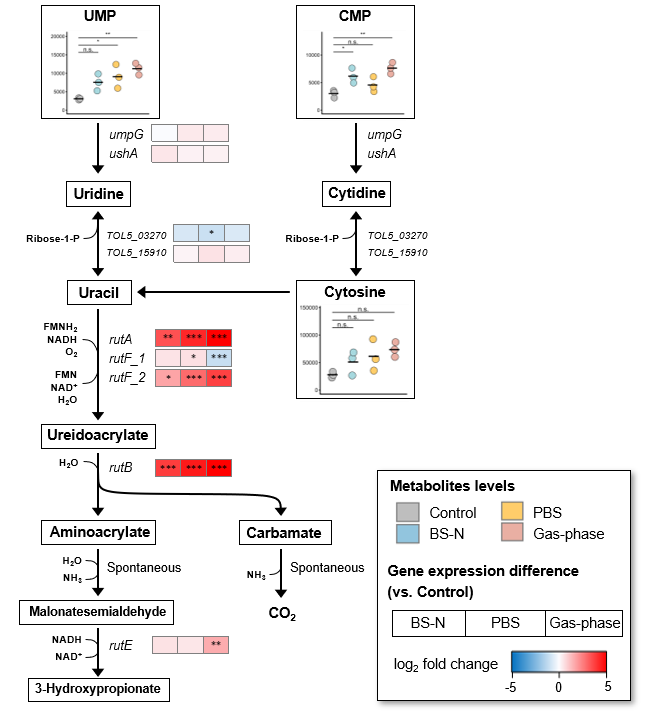

**Figure S11.** Metabolite levels and differential gene expression for pyrimidine metabolism. For metabolites, each dot represents the normalized peak height of one of three biological replicates from the control, BS-N, PBS, and gas-phase conditions, and the bars indicate the mean values. Asterisks indicate statistical significance based on adjusted *p* values obtained by Dunnett's test (* *p* < 0.05, ** *p* < 0.01, *** *p* < 0.001, **** *p* < 0.0001), with the control group as the reference. For gene expression, the boxes represent the log_2_ fold change of gene expression levels in the BS-N, PBS, and gas-phase conditions relative to the control. The color gradient indicates the expression difference. Asterisks within the boxes indicate significance derived from the false discovery rate (FDR) calculated in the differential gene expression analysis (* FDR < 0.05, ** FDR < 0.01, *** FDR < 0.001). Abbreviations: UMP, uridine 5′-monophosphate; CMP, cytidine 5′-monophosphate; Ribose-1-P, α-D-ribose 1-phosphate.

**Supplemental tables**

**Table S1.** Details of the internal standards for hydrophilic metabolomics. (Excel file)

**Table S2.** Details of the internal standards for lipidomics. (Excel file)

**Table S3.** Information about the pairs of internal standards and lipid subclasses. (Excel file)

**Table S4.** Differential expressed genes in Tol 5 under the BS-N, PBS, Gas-phase condition. (Excel file)

**Table S5.** Differential gene expression of Tol 5 cells in the BS-N, PBS, and gas-phase conditions relative to those in the control condition.

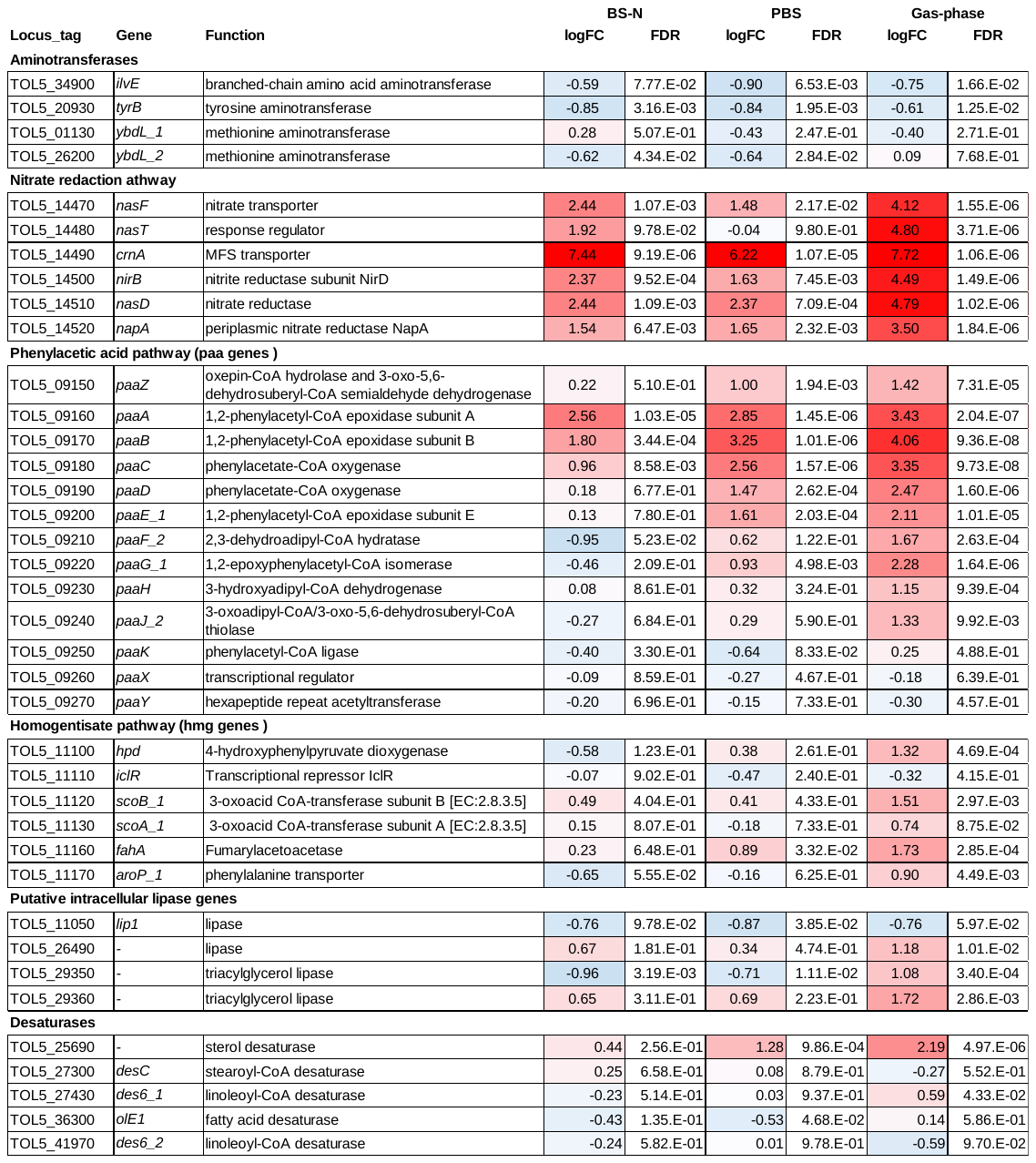

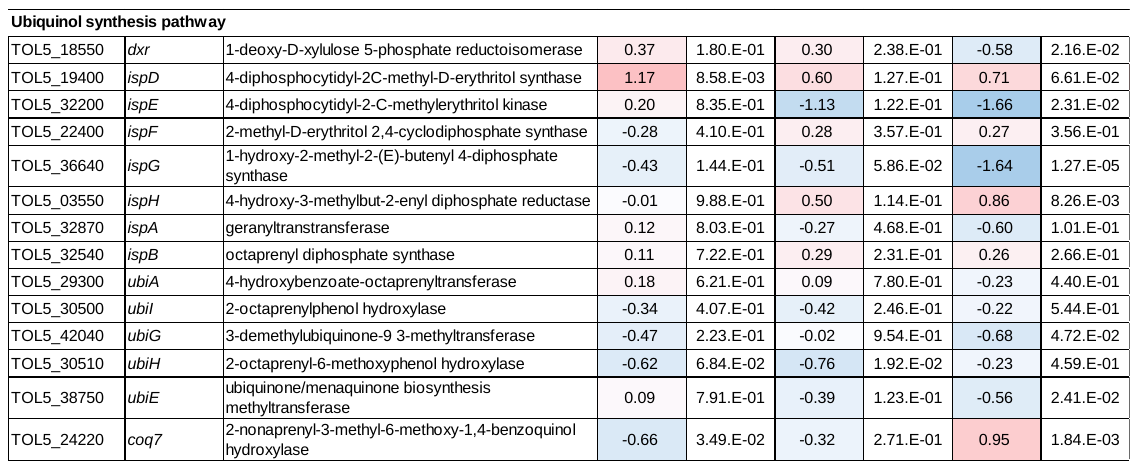
